# Structural and molecular determinants of CCS-mediated copper activation of MEK1/2

**DOI:** 10.1101/2020.05.01.072124

**Authors:** Michael Grasso, Gavin J. Bond, Ye-Jin Kim, Katherine B. Alwan, Stefanie Boyd, Maria Matson Dzebo, Sebastian Valenzuela, Tiffany Tsang, Natalie A. Schibrowsky, Megan L. Matthews, George M. Burslem, Pernilla Wittung-Stafshede, Duane D. Winkler, Ninian J. Blackburn, Ronen Marmorstein, Donita C. Brady

**Affiliations:** Department of Chemistry, School of Arts and Sciences, University of Pennsylvania, Philadelphia, PA, 19104, USA; Abramson Family Cancer Research Institute, Perelman School of Medicine, University of Pennsylvania, Philadelphia, PA, 19104, USA; Biochemistry Major Program, College of Arts and Sciences, University of Pennsylvania, Philadelphia, PA, 19104, USA; Department of Cancer Biology, Perelman School of Medicine, University of Pennsylvania, Philadelphia, PA, 19104, USA; Department of Chemical Physiology and Biochemistry, School of Medicine, Oregon Health and Science University, Portland, OR, 97239, USA; Department of Biological Sciences, University of Texas at Dallas, Richardson, TX, 75080, USA; Department of Biology and Biological Engineering, Chalmers University of Technology, 412 96 Gothenburg, Sweden; Cell and Molecular Biology Graduate Group, Perelman School of Medicine, University of Pennsylvania, Philadelphia, PA, 19104, USA; Department of Biochemistry and Molecular Biophysics, Perelman School of Medicine, University of Pennsylvania, Philadelphia, PA, 19104, USA

**Keywords:** Transition metal, Protein Kinase, Signal Transduction, X-ray Crystallography, Chemical Biology

## Abstract

Normal physiology relies on the precise coordination of intracellular signal transduction pathways that respond to nutrient availability to balance cell growth and cell death. We recently established a critical mechanistic function for the redox-active micronutrient copper (Cu) in the canonical mitogen activated protein kinase (MAPK) pathway at the level of MEK1 and MEK2. Here we report the X-ray crystal structure of Cu-MEK1 and reveal active site chemical ligands and oxidation state specificity for MEK1 Cu coordination. Mechanistically, the Cu chaperone CCS selectively bound to and facilitated Cu transfer to MEK1. Mutations in MEK1 that disrupt Cu(I) affinity or a CCS small molecule inhibitor reduced Cu-stimulated MEK1 kinase activity. These atomic and molecular level data provide the first mechanistic insights of Cu kinase signaling and could be exploited for the development of novel MEK1/2 inhibitors that either target the Cu structural interface or blunt dedicated Cu delivery mechanisms via CCS.

## Introduction

The development of a multicellular organism depends on the precise synchronization of cell division, death, specialization, interactions, and movement. The orchestration of complex cellular processes in response to environmental changes is essential for organismal tissue homeostasis. Intrinsic molecular mechanisms enable conversion of fluctuating extracellular and intracellular inputs into various outputs that drive cellular processes necessary for adaptation(Efeyan et al., 2015; Kolch et al., 2015). This interpretation and integration of inputs is controlled by signal transduction pathways that facilitate the precise coordination of cellular processes with dynamic spatial and temporal control(Efeyan et al., 2015; Kolch et al., 2015). One of the simplest evolutionarily conserved mediators of signal transduction is the protein kinase, an enzyme that traditionally phosphorylates a substrate at threonine, serine, and/or tyrosine residues(Manning et al., 2002). Kinases within complex protein modules directly respond to and/or sense growth factors, nutrients, and metabolites to influence signal amplification, duration, and frequency(Efeyan et al., 2015; Kolch et al., 2015).

While the kinase signaling networks that integrate fluctuations in the abundance of many nutrients and metabolites are well-established(Efeyan et al., 2015; Kolch et al., 2015), the discovery of signaling molecules capable of mediating similar functions downstream of transition metal availability is underdeveloped. Traditionally, the redox-active transition metal copper (Cu) functions as a high affinity structural and/or catalytic cofactor within the active site of Cu-dependent enzymes(Chang, 2015; Kim et al., 2008; Thiele and Gitlin, 2008). Phenotypic studies of the rare genetic diseases, Menkes and Wilson, solidified the physiological impact of aberrant Cu absorption and excretion(Ala et al., 2007; Kaler, 2014; Mercer et al., 1993; Yamaguchi et al., 1993). Mechanistically, these diseases display deficiencies in cellular functions attributed to the dozens of known Cu-dependent enzymes and helped elucidate the cellular machinery responsible for the proper acquisition and distribution of Cu(Ala et al., 2007; Kaler, 2014; Kim et al., 2008). Despite the need for strict homeostatic mechanisms to control Cu abundance, we know relatively little about direct kinase signaling pathways that respond to and or/sense Cu abundance, especially those that integrate to influence cellular proliferation.

In pursuit of providing a molecular explanation for the observations that biological systems convert Cu abundance into cellular decisions like proliferation(Turski and Thiele, 2009), we recently reported an unexpected link between the cellular acquisition of Cu and a mitogenic signaling cascade instrumental in cell proliferation(Turski et al., 2012). Namely, genetic ablation of the primary Cu transporter gene *Ctr1*, which in multiple organisms results in growth retardation and embryonic lethality(Dancis et al., 1994; Lee et al., 2002; Turski and Thiele, 2007), reduced mitogen activated protein kinase (MAPK) signaling downstream of growth factor stimulated receptor tyrosine kinase (RTK) activation(Turski et al., 2012). The canonical MAPK pathway consists of the RAF-MEK-ERK signaling cascade and represents one of the most well-defined axes within eukaryotic cells to promote cell proliferation(Morrison, 2012). In response to RTK activation, GTP-bound RAS engages the serine/threonine RAF kinases via their RAS binding domains (RBD) to promote phosphorylation and activation of the MEK1 and MEK2 kinases, which in turn phosphorylate and activate the ERK1 and ERK2 kinases to drive cell proliferation(Cobb, 1999; Terrell and Morrison, 2019). Layered on top of this signaling cascade are multiple positive and negative regulatory mechanisms that fine-tune the frequency, duration, localization, and amplitude of canonical MAPK signaling(Friedman and Perrimon, 2006a), underscoring the importance of tightly regulating this pathway at the cellular and organismal level for a host of responses. Genetic, molecular, and biochemical interrogation of the requirement of Cu for MAPK pathway activation revealed a direct metal-kinase interaction between Cu and MEK1/2 that was sufficient to augment phosphorylation and activation of the ERK1 and ERK2 kinases(Turski et al., 2012). This is the first example of Cu directly regulating the activity of a mammalian kinase and has exposed a signaling paradigm that directly connects Cu to signaling pathway components(Chang, 2015). Further, targeting aberrant MAPK signaling in melanomas and other cancers(Caunt et al., 2015; Davies et al., 2002; Holderfield et al., 2014) contributes to the efficacy of Cu chelators(Brady et al., 2014, 2017; Kim et al., 2020; Xu et al., 2018) that are traditionally utilized to treat Cu overload in Wilson disease patients(Ala et al., 2007) and highlights the importance of dissecting the molecular mechanism of MEK1/2 activation via Cu.

To distinguish the pleiotropic impact of global reductions in Cu transport from the requirement of Cu for MEK1/2 kinase activity and MAPK signaling, we previously undertook targeted mutagenesis and metal-catalyzed oxidation reactions followed by mass spectrometry to identify a mutant of MEK1 (M187A, H188A, M230A, H239A) devoid of Cu-binding and kinase activity(Brady et al., 2014). Despite this previous biochemical analyses, the precise atomic level interaction and the mechanism of activation has yet to be determined (Brady et al., 2014). In the current study, we set out to systematically interrogate the structural and molecular determinants of MEK1 Cu-binding and provide a mechanism by which selective Cu delivery and subsequent MEK1/2 activation is achieved. We report the X-ray crystal structure of Cu bound MEK1, which was corroborated in-solution by XAS and EXAFS studies of Cu(I)-MEK1. *In vitro* and in mammalian cells Cu-stimulated MEK1/2 kinase activity required the Cu chaperone for superoxide dismutase (CCS) and could be blocked with the small molecule inhibitor DCAC50 that targets CCS. Taken together, the data provide a more precise understanding of how Cu cooperates with MEK1/2 that can be utilized to expand the landscape of Cu in kinase signal transduction.

## Results

### The X-ray crystal structure of MEK1 with Cu reveals active site chemical ligands

We first investigated the Cu binding mechanism of MEK1 at the atomic level by producing a previously crystallized, soluble MEK1 fragment in *Escherichia coli*(Meier et al., 2012). The MEK1 fragment consisted of amino acids 45-393 with a flexible linker region between amino acids 264-307 replaced with a 6-residue SVGSDI linker(Meier et al., 2012). This MEK1 protein construct was subsequently purified to homogeneity and crystallized in the space group P 1 2_1_ 1 with two molecules of MEK1 within the asymmetric unit to a resolution of 2.3Å (Table 1). Despite the asymmetric unit containing two MEK1 molecules (Figure 1A), this interaction is thought to be an artifact of crystal packing due to the quaternary structure varying from what was previously reported to be a biologically relevant dimer(Ohren et al., 2004). While crystals were formed in the presence of magnesium chloride (MgCl_2_) and AMP-PNP, a non-hydrolysable ATP analog, Mg is absent in the structure. Also, the AMP-PNP molecules have unresolved γ-phosphates, resulting in the need to model in ADP in the MEK1 subunits (Figure 1A). We utilized structures derived from these crystals to interrogate the mechanism of MEK1 Cu binding.

**Table 1.** Summary of crystallographic statistics.

| **Crystal** | **MEK<sup>45-393</sup>/SVQSDI** | **MEK<sup>45-393</sup>/SVQSDI/Cu** |
| --- | --- | --- |
| **Resolution Range (Å)** | 47.67 - 2.257 (2.338 - 2.257) | 41.96 - 2.381 (2.466 - 2.381) |
| **Space Group** | P 1 2<sub>1</sub> 1 | P 1 2<sub>1</sub> 1 |
| **Unit Cell (a, b, c, α, β, γ)** | 46.65 95.35 72.81 90 93.02 90 | 46.88, 95.04, 72.18, 90, 94.13, 90 |
| **Total Reflections** | 324593 | 762036 |
| **Unique Reflections** | 29319 (2864) | 24847 (2447) |
| **Redundancy** | 2.9 (2.7) | 4.6 (3.6) |
| **Completeness (%)** | 98.1 (97.9) | 98.00 (96.49) |
| **Mean I/sigma (I)** | 17.63 (2.66) | 23.25 (3.19) |
| **Wilson B Factor** | 33.91 | 35.81 |
| **R-merge** | 0.118 (0.421) | 0.147 (0.484) |
| **R-meas** | 0.143 (0.512) | 0.165 (0.560) |
| **R-pim** | 0.080 (0.287) | 0.074 (0.275) |
| **CC1/2** | 0.982 (0.699) | 0.998 (0.821) |
| **Reflections used in refinement** | 29314 (2863) | 24842 (2445) |
| **Reflections used for R-free** | 1989 (191) | 2008 (204) |
| **R-work** | 0.1816 (0.2303) | 0.1900 (0.2482) |
| **R-free** | 0.2334 (0.3127) | 0.2479 (0.3221) |
| **RMS (bonds)** | 0.009 | 0.006 |
| **RMS (angles)** | 1.32 | 0.85 |
| **Ramachandran Favored (%)** | 97.64 | 97.73 |
| **Ramachandran Outliers (%)** | 0.17 | 0.00 |
| **Rotamer Outliers** | 1.02 | 0.64 |
| **Clashscore** | 10.17 | 12.83 |
| **Average B Factor** | 39.74 | 42.58 |

**Figure 1.**
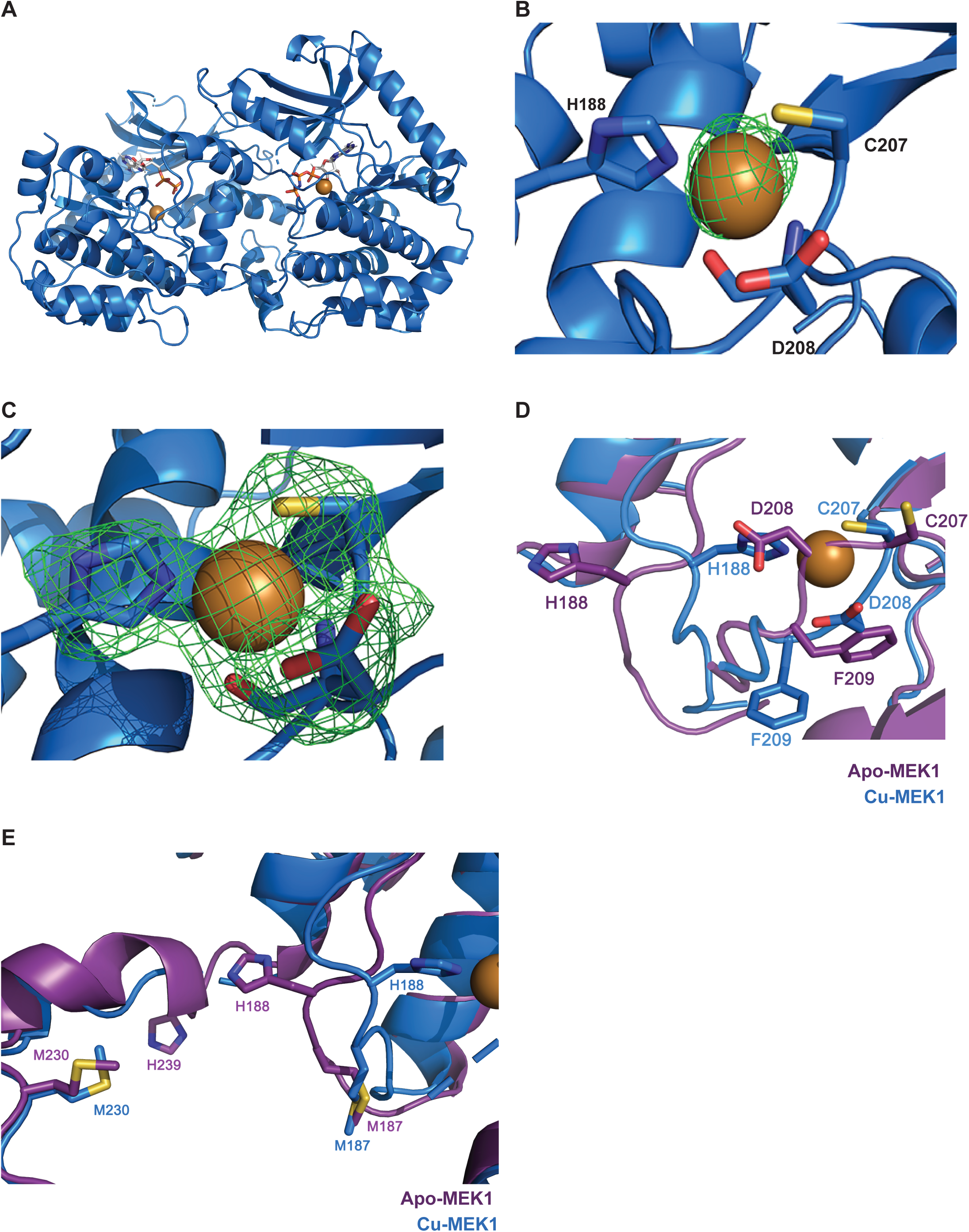
X-ray crystal structure of Cu bound MEK1. (A) Crystal structure of MEK1^45-393/SVGSDI^ molecules (marine blue) bound with Cu ions (brown) and AMP-PNP (grey). (B) Close-up view of Cu ion (brown) with its anomalous map (lime green, contoured at 3.0 sigma) bound to MEK1 (marine blue) at the sidechains of H188 and C207 and the main chain of D208. (C) Close-up view of Cu ion (brown) and MEK1 interactions at H188, C207, and D208 with their simulated annealing omit map (lime green, contoured at 3.0 sigma). (D) Close-up view of alignment of crystal structures of apo MEK1 MEK1^45-393/SVGSDI^ (purple) bound with AMP-PNP and MEK1^45-393/SVGSDI^ (marine blue) bound with Cu ions (brown) and AMP-PNP with Cu interacting sidechains H188, C207, and D208 indicated and F210 of DGF motif. (E) Structural proximity between proposed Cu “entry site” containing M187, M230, and H239 and Cu active site containing H188 of the apo MEK1 MEK1^45-393/SVGSDI^ (purple) bound with AMP-PNP and MEK1^45-393/SVGSDI^ (marine blue) bound with Cu ions and AMP-PNP.

The Cu-MEK1 crystal structure was obtained by soaking crystals in 2-fold molar excess cupric chloride (CuCl_2_) and then collecting data at the anomalous edge of Cu (9000 eV) to ensure that the Cu binding site could be unambiguously identified. Interrogation of the anomalous map contoured to 3 sigma revealed a substantial signal near residues histidine 188 (H188), cysteine 207 (C207), and aspartic acid 208 (D208) (Figure 1B). Cu coordinated with the nitrogen of H188 (2.1Å), the sulfur atom of C207 (2.4Å), and the carbonyl and the amide nitrogen of D208 (2.2Å and 2.1Å, respectively) (Figure 1B). This coordination forms a square planar arrangement of the Cu atom between all four amino acid atoms, which is characteristic of Cu(II) binding (Figure 1C). Interestingly, the sidechain of D208 “caps” the Cu by protecting the top of the atom at a distance of 3.5-3.7 Å, far enough away to not be considered coordinated (Figure 1C). While Mg is not visible in either crystal structure, the D208 sidechain necessary for Mg coordination is not a ligand for the Cu atom (Figure 1C). Interestingly, both the difference map (f_o_-f_c_) and anomalous map indicate a small amount of unresolved density above the main density where Cu is modeled in chain A (Figures 1A and S1A). The density is not large enough to occupy a separate atom, so two conformers were modelled of the Cu atom in chain A, with the second density demonstrating 20% occupancy (Figure S1A). In this position, the Cu atom maintains contact with C207 but not H188 (Figure S1B). Therefore, this second occupancy could indicate an intermediate Cu-MEK1 species and/or a level of flexibility allowed for the Cu atom within its canonical binding mode (Figures S1A and S1B).

Since Cu binds within the MEK1 active site, we next investigated whether the binding confers conformational changes. The RMSD between the apo- and Cu-MEK1 structures was negligible at 1.447Å. However, Cu introduction resulted in extensive changes in residues 183-190, as well as the DFG motif, which lies directly after C207 (Figures 1D and S1C). H188 of the R1 position of the regulatory spine flips in dramatically from its typical anchored position to aspartate 245 (D245) of the F-helix by a hydrogen bond, resulting in large changes in residues 183-190 (Figures 1D and S1C). C207 moves inward to coordinate the Cu atom and D208 moves outward to allow its main chain atoms to coordinate while the sidechain “caps” the Cu (Figures 1D and S1C). This movement results in significant displacement of F209, but the rest of the activation segment (residues 211-233) aligns with the unbound structure (Figures 1D and S1C). Despite this movement, both structures display a “DFG-in” conformation (Figures 1D and S1C). Interestingly, the residues immediately following the activation segment (236-241) are disordered and unresolved in the Cu-MEK1 structure but resolved in the apo-MEK1 structure, indicating that Cu binding may also affect this region of the kinase (Figures 1D and S1C). Of note, methionine 230 (M230) and histidine 239 (H239) adjacent to and within this disordered, unresolved region were previously reported to contribute to MEK1 Cu binding and kinase activity(Brady et al., 2014), Specifically, mutation of M187, H188, M230, or H239, which are conserved in MEK2 that also binds Cu and is inhibited by the Cu chelator tetrathiomolybdate, TTM, alone or in combination to alanine abrogated the ability of MEK1 to bind a Cu-charged resin and phosphorylate ERK1/2 with undetectable impact on protein folding(Brady et al., 2014). We hypothesize that Cu may either interact with MEK1 at these two distinct sites, M198/M230/H239 and H188/C207/D208, which induces a dynamic conformational change that contributes to MEK1 activation, or binds to an “entry site” consisting of M187/M230/H239 prior to being transferred to the active site, H188/C207/D208 (Figure 1E). Taken together, these data provide the first evidence of Cu coordination by a mammalian protein kinase, along with the necessary chemical ligands that confer changes in protein structure.

### MEK1 Cu binding is oxidation state selective

We next confirmed MEK1 Cu binding capability in-solution by measuring Cu to protein stoichiometry by inductively coupled plasma mass spectrometry (ICP-MS). Cu(II) ions were introduced as cupric sulfate (CuSO_4_) to recombinant, full length MEK1 at 2.5 molar excess or stoichiometric levels (1:1), followed by removal of excess. Both reconstitutions resulted in an average of 0.5 Cu atoms bound per MEK1 protomer (Table 2). To determine whether Cu was bound as Cu(II) or Cu(I), electron paramagnetic resonance (EPR) spectra of Cu(II) MEK1 and a series Cu(II)-EDTA standards were collected (Figure 2A; Table 2). The EPR spectrum of the Cu(II) MEK1 was devoid of signal, indicating that the Cu was bound in an EPR undetectable state (Figure 2A; Table 2). These results exclude mononuclear Cu(II) binding, and strongly suggest a reductive process which can generate Cu(I) *in situ*, which subsequently binds to cuprophilic sites in MEK1. However, other scenarios such as antiferromagnetically coupled Cu(II) dimeric structures cannot be ruled out on the basis of the EPR data alone.

**Table 2.** Cu(II) binding and EPR properties of Cu(II) reconstituted MEK1.

| **Sample** | **Protein**<br>**($\mu\text{M}$)** | **Cu($\mu\text{M}$)** | **Cu/**<br>**protein** | **Integral/**<br>**$10^4$** | **Integral/$10^4$**<br>**corrected**<br>**for buffer** | **EPR**<br>**detectable**<br>**Cu ($\mu\text{M}$)** | **% EPR**<br>**detectable**<br>**Cu** |
| --- | --- | --- | --- | --- | --- | --- | --- |
| **MEK1<sup>WT</sup> #1** | 290 | 152 | 0.52 | ND | ND | ND | ND |
| **MEK1<sup>WT</sup> #2** | 380 | 209 | 0.55 | 43.3 | -0.86 | -2.1 | -0.5 |
| **CuEDTA #1** |  | 104 |  | 403.8 | 359.6 | 104 | 100 |
| **CuEDTA #2** |  | 216 |  | 1120.6 | 1076.4 | 216 | 100 |
| **CuEDTA #3** |  | 323 |  | 1727.4 | 1285.6 | 323 | 100 |
| **CuEDTA #4** |  | 421 |  | 2200.6 | 2156.4 | 421 | 100 |
| **Buffer (50**<br>**mM**<br>**HEPES)** | 0 | 0 |  | 44.1 | 0 | 0 | 0 |

**Figure 2.**
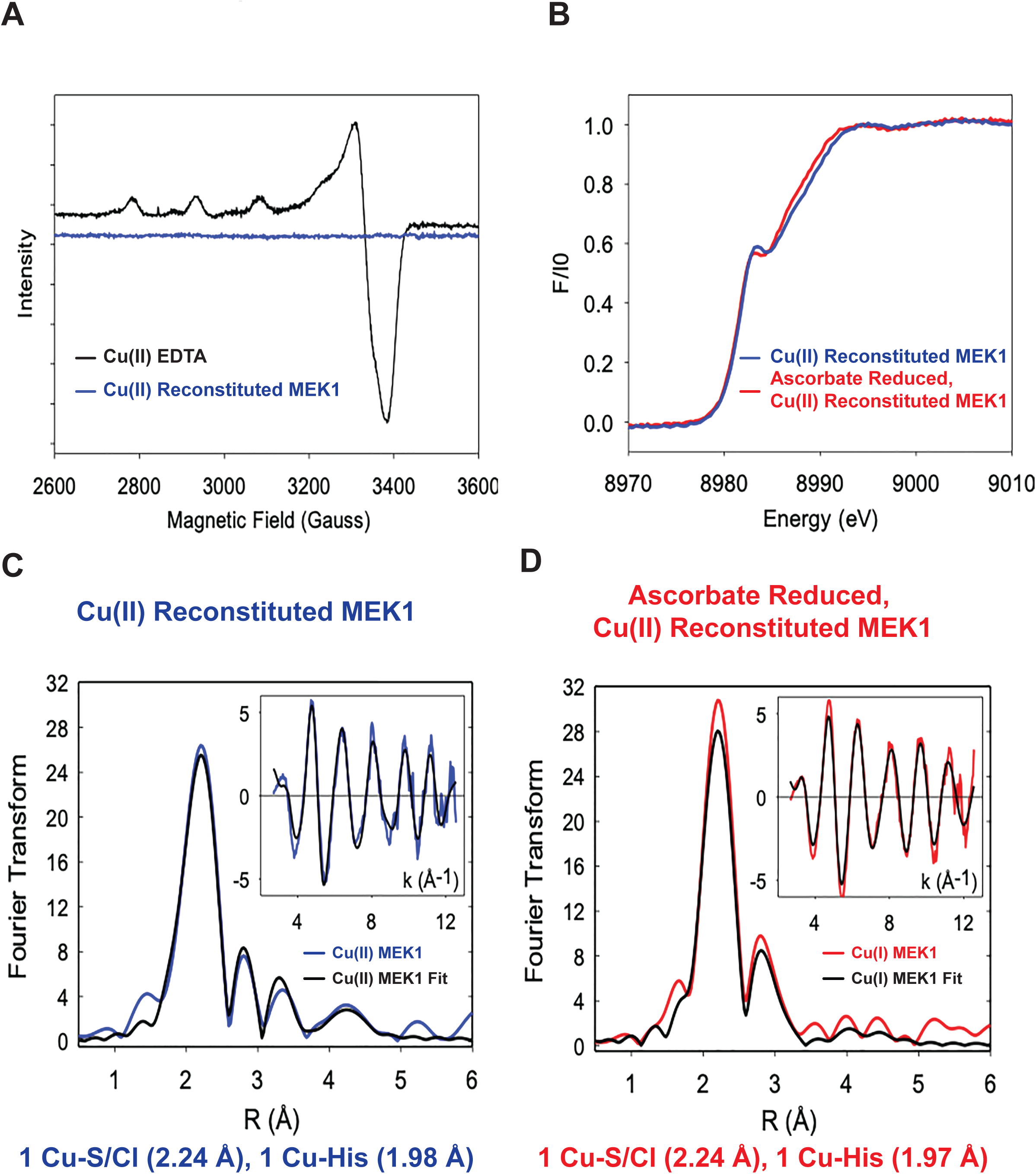
MEK1 binds Cu(I) at high affinity via histidine and cysteine ligands. (A) EPR spectra of Cu(II) reconstituted MEK1 (blue line) or Cu(II) EDTA standard (black line). n=2 technical replicates. (B) XAS of Cu K absorption edges of Cu(II) reconstituted MEK1 (blue line) or ascorbate reduced, Cu(II) reconstituted MEK1 (red line). n=2 technical replicates. (C and D) Fourier transform and EXAFS (inset) of (C) experimental Cu(II) reconstituted MEK1 (blue line) in NaCl and its fit (black line) or (D) experimental ascorbate reduced, Cu(II) reconstituted MEK1 in NaCl (red line) and its fit (black line). n=2 technical replicates.

### MEK1 binds Cu(I) at high affinity via nitrogen and sulfur ligands

To probe further the oxidation state and coordinate structure of Cu bound MEK1 in-solution, X-ray absorption spectroscopy (XAS) experiments were conducted on two MEK1 samples: (i) the product of Cu(II) reconstitution and (ii) after ascorbate reduction (Figure 2B). Based on the EPR spectra (Figure 2A), we hypothesized that the samples would be identical if simple reductive chemistry is responsible for generating the Cu(I) state from Cu(II). Comparison of the absorption edges of the Cu(II)-reconstituted MEK1 and ascorbate reduced, Cu(II)-reconstituted MEK1 indicated very close correspondence between the spectra, as further evidence that a similar Cu(I)-bound state is achieved in each case (Figure 2B). Extra X-ray absorption fine structure (EXAFS) spectra of the two MEK1 samples, confirmed these similarities (Figures 2C and 2D) and are indicative of 1 Cu-S/Cl interaction at an average distance of 2.24Å and 1 His residue at 1.98-2.01Å (Figures 2C and 2D; Table 3). Both Cu(II)-reconstituted and ascorbate-reduced MEK1 spectra are better simulated by inclusion of two additional shells, an O/N at ∼2.5Å and a S/Cl scatterer at ∼2.8Å (Table 3). In agreement with the Cu-MEK1 crystal structure, which captured a Cu(II)-bound species that exists prior to MEK1 oxidation and subsequent Cu reduction that occurs *in situ*, these overlapping in-solution findings indicate that the coordination is consistent with a Cu(I) center depicted as binding to H188, C207, and D208, but with only weak association of ligands from the D208 residue together with a weak association with a halide from the buffer. To gain further information on the contribution of Cl to the 2.24Å or for the 2.85Å scatterer, Cl ions were substituted in the buffer with 150 mM KBr. XAS and EXAFS of the bromide-treated ascorbate-reduced, Cu(II)-reconstituted MEK1 could be simulated by the same ligand set as before except for Cu-Br replacing the 2.8Å Cu-Cl now at a closer distance of 2.53Å (Figure S2A; Table 3). The decrease in Cu-halide distance is significant since the increased covalency between the Cu(I) center and the more polarizable bromide ligand is expected to decrease the Cu to bromide distance. Finally, bicinchoninic acid (BCA) ligand competition experiments were performed with MEK1^WT^ and a point mutant at C207 to alanine to obtain precise Cu(I) binding affinity. While MEK1^WT^ binds Cu(I) at an affinity of 9.24 ± 3.34 ^E-16^M(Xiao et al., 2011), mutation at C207 significantly reduced Cu(I) affinity (Table 4), suggesting that C207 identified in the Cu(II) X-ray crystal structure is necessary for Cu(I) binding. Collectively, these findings indicate MEK1 coordinates Cu(I) via histidine and sulfur ligands and suggest that thiol-disulfide redox chemistry may contribute to the observed *in situ* reduction of the Cu ion.

**Table 3.**
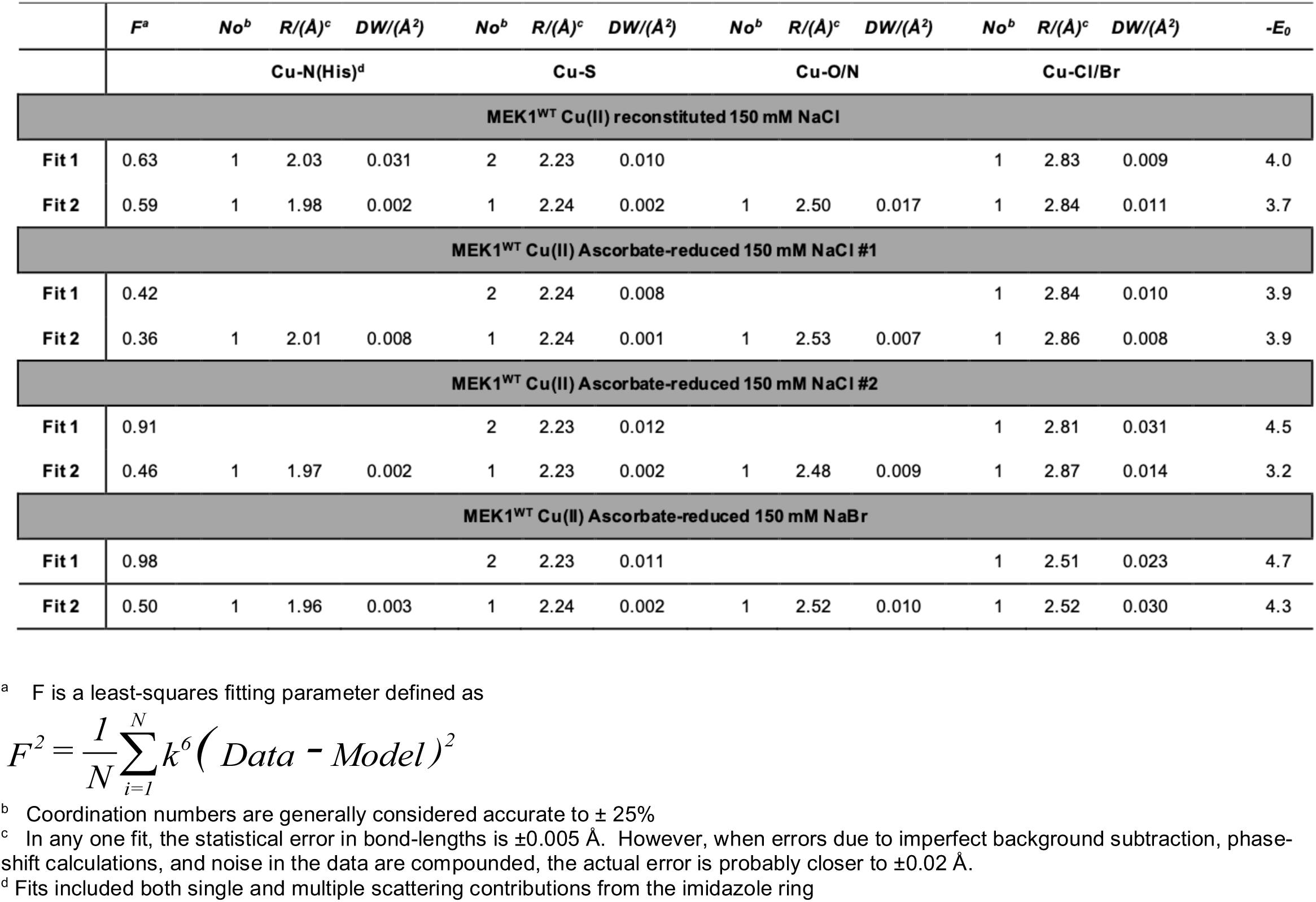
Parameters associated with simulation of the EXAFS spectra of MEK1 samples using EXCURVE 9.2.

**Table 4.** Cu(I) Dissociation Constant for MEK1.

| **Protein** | ***K<sub>d</sub>*** |
| --- | --- |
| **MEK1<sup>WT</sup>** | 9.24 +/- 3.34 E<sup>-16</sup> M |
| **MEK1<sup>C207A</sup>** | < E<sup>-11</sup> M |

### Mutation of the MEK1 Cu binding ligands reduces MEK1 kinase activity

We next investigated mutants at H188 and C207 of MEK1, based on the crystal structure and in-solution studies, in an ELISA-based *in vitro* kinase assay. Both MEK^H188A^ and MEK^C207A^, which are likely devoid of any Cu as isolated, had negligible detectable kinase activity when compared to MEK1^WT^ that binds between 0.1 and 0.5 Cu atoms per MEK1 protomer(Brady et al., 2014; Turski et al., 2012) (Figure 3A). Previous targeted mutagenesis of MEK1 amino acids conserved between *Drosophila* and human MEK1 and are known to coordinate Cu, including H188 and C207, resulted in similar reductions in kinase activity in mammalian cells(Brady et al., 2014). Since H188 is part of the HRD domain, which functions in catalytic activity, and C207 lies directly in front of the DFG motif, which coordinates Mg and positions the activation loop(Roskoski, 2012), mutations that could weakly coordinate Cu were introduced to evaluate whether MEK1 activity could be rescued. However, MEK1^H188M^ and MEK1^C207S^ retained minimal ability to phosphorylate ERK2 (Figure 3A). Further, substituting H188 for phenylalanine in the HRD motif, chosen because this amino acid does not bind Cu and more closely resembles the histidine side chain, reduced MEK1 kinase activity (Figure 3A), suggesting a requirement for Cu binding, and not changes to HRD-mediated stabilization of the kinase regulatory spine(McClendon et al., 2014; Meharena et al., 2013) (Figure 3A).

**Figure 3.**
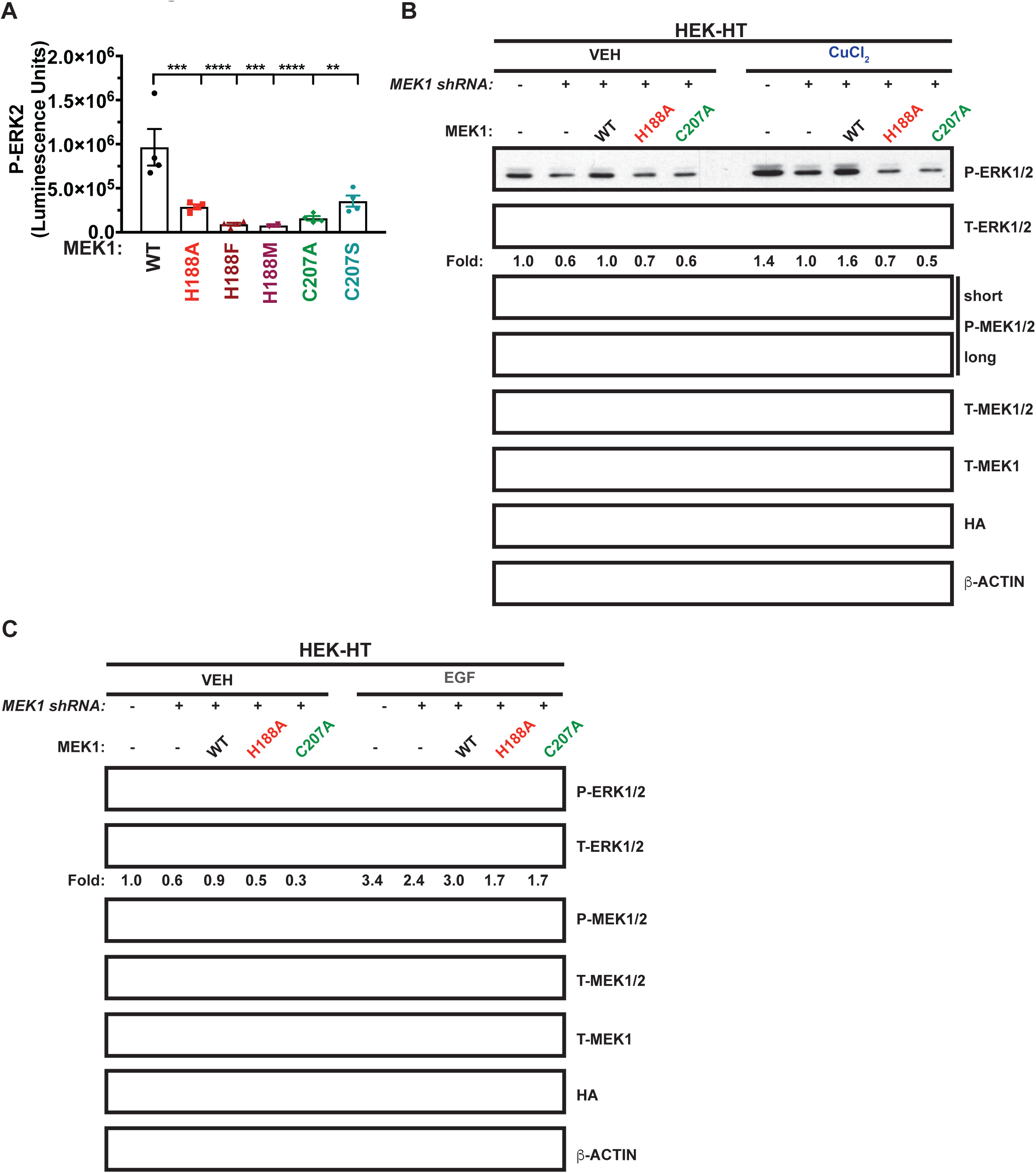
MEK1 Cu binding is required for kinase activity. (A) Scatter dot plot with bar at mean luminescence units ± s.e.m. from ELISA detection of recombinant phosphorylated (P)-ERK2^K54R^ from MEK1^WT^, MEK1^H188A^(H188A), MEK1^H188F^(H188F), MEK1^H188M^(H188M), MEK1^C207A^(C207A), or MEK1^C207S^(C207S). n=4 technical replicates or MEK1^H188M^ n=2 technical replicates. Results were compared using a one-way ANOVA followed by a Dunnett’s multi-comparisons test. **, P=0.0015; ***; P=0.0006; ****, P<0.0001. (B and C) Immunoblot detection of P-ERK1/2, total (T)-ERK1/2, P-MEK1/2, T-MEK1/2, T-MEK1, HA, or β-ACTIN from HEK-HT cells stably expressing *shRNA* against control (-) or *MEK1* reconstituted with *HA-MEK1*^*WT*^(WT), *HA-MEK1*^*H188A*^ (H188A), or *MEK1*^*C207A*^(C207A) treated with (B) vehicle (VEH) or 1µM CuCl_2_ for 20 minutes or (C) VEH or 0.1ng/mL EGF for 15 minutes. Quantification: Fold change P-ERK1/2/T-ERK1/2 normalized to control, VEH. n=3 biologically independent experiments.

To evaluate the consequence of disrupting MEK1 Cu-binding on MAPK signaling, *MEK1* knockdown immortalized human embryonic kidney cells (HEK-HT) were generated via short hairpin (shRNA) and validated to have reduced MEK1 protein expression and as a consequence diminished phosphorylation of ERK1/2 (P-ERK1/2) (Figures 3B and 3C). Interestingly, CuCl_2_ addition to HEK-HT cells for only 20 minutes was sufficient to selectively elevate P-ERK1/2 but not phosphorylation of MEK1/2 (Figures 3B and 3C), indicating that MEK1/2 are not saturated with Cu in cells and supporting the idea that Cu is a bonafide mediator of MAPK signaling. These kinetics of MEK1/2 activation are consistent with rapid cellular uptake of Cu via CTR1(Maryon et al., 2013). Further, we tested and found that the decrease in P-ERK1/2 in HEK-HT cells due to knockdown of endogenous *MEK1* either basally or in response to CuCl_2_ or EGF was rescued by expressing wild-type but not H188A and C207A mutants of MEK1 (Figures 3B and 3C). Thus, mutations at the ligands essential for Cu binding to MEK1 reduced MAPK signaling, bolstering the importance of Cu as both an activator and modulator of the pathway.

### CCS binds MEK1 to facilitate Cu binding and increased kinase activity

We next explored the existence of a dedicated cellular Cu delivery mechanism capable of supporting Cu-stimulated MEK1/2 activity (Figure 3B). Evolutionarily conserved Cu chaperones facilitate the efficient delivery of Cu to cuproenzymes located in the cytosol, trans-Golgi network, or mitochondria(Robinson and Winge, 2010). Based on the predominant cytosolic localization of MEK1/2, we tested whether the cytosolic Cu chaperones ATOX1 or CCS bind to MEK1 in single-cycle SPR experiments (Figures 4A and 4B). Recombinant MEK1 was coupled to a SPR sensor followed by injection of increasing concentrations of PARVALBUMIN (Figure S3A), as a negative control, or the Cu chaperones ATOX1 or CCS (Figures 4A and 4B). CCS interacted with MEK1 at an apparent dissociation constant (K_D_) of 2.09±0.48µM (Figure 4B; Table 5) when compared to ATOX1 (Kd=10.9±5.65µM), confirming a direct interaction between CCS and MEK1. However, both EMSA assays and size exclusion chromatography (SEC) of excess MEK1 or CCS indicate that the association between apo-CCS or Cu-CCS and MEK1 is transient based on the absence of a stable complex that could be co-eluted (Figures S3B-S3E).

**Table 5.** Association (k_a_), dissociation rate constant (k_d_), dissociation constant (K_d_), and predicted maximum response (R_max_) of ATOX1 or CCS binding to MEK1 obtained with SPR.

| | $k_a$ ($M^{-1}s^{-1}$) | $k_d$ ($s^{-1}$) | $K_d$ ($\mu M$) | $R_{max}$ (RU) |
| --- | --- | --- | --- | --- |
| **ATOX1** | 309±109 | $(3.07 \pm 0.55) \cdot 10^{-3}$ | 10.9±5.65 | 28.67±3.92 |
| **CCS** | 533±163 | $(1.06 \pm 0.12) \cdot 10^{-3}$ | 2.09±0.48 | 67.39±4.92 |

SPR data was fitted with a 1:1 binding model and the presented values are the average and standard deviation of mean ( $n=3$ independent experiments).

**Figure 4.**
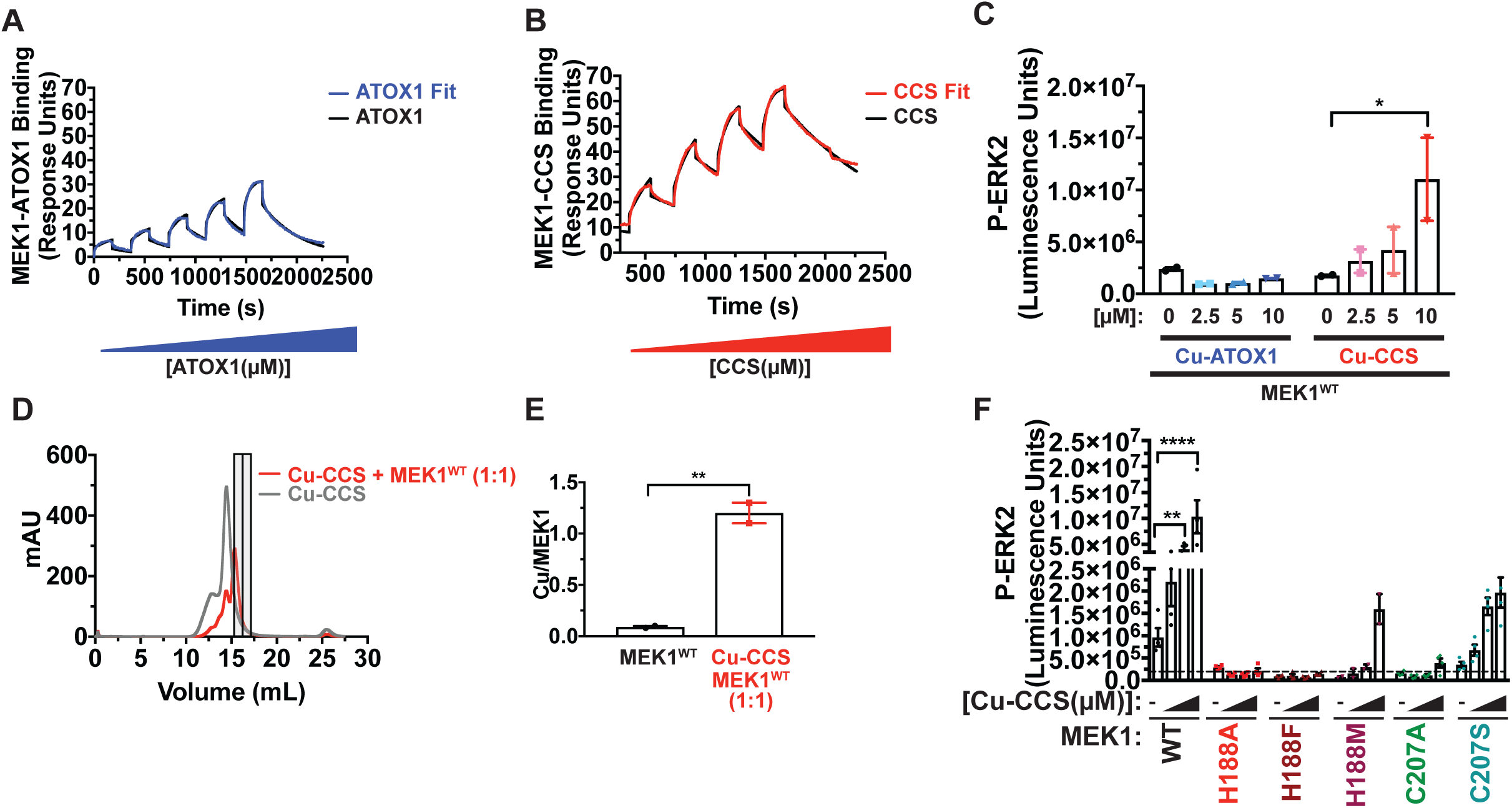
CCS directly interacts with MEK1 to facilitate Cu binding and kinase activity. (A and B) Single-cycle SPR experiments of ATOX1 or CCS binding to MEK1 in which increasing concentrations of (A), ATOX1 (2.5-40 µM, blue line) or (B), CCS (5-40 µM, red line) was injected onto immobilized MEK1 and their 1:1 binding model fits (black lines). n=3 independent experiments. (C) Scatter dot plot with bar at mean luminescence units ± s.e.m. from ELISA detection of recombinant phosphorylated (P)-ERK2^K54R^ from MEK1^WT^ incubated with increasing concentrations of Cu reconstituted ATOX1 (Cu-ATOX1) or CCS (Cu-CCS). n=2 technical replicates. Results were compared using a two-way ANOVA followed by a Tukey’s multi-comparisons test. *, P=0.0185. (D) Size exclusion chromatography runs of Cu-CCS (grey line) or Cu-CCS and MEK1 mixed 1:1 (red line). Bars indicated MEK1 positive fractions 15 and 16 from Cu-CCS and MEK1 mixed 1:1 analyzed by inductively coupled plasma mass spectrometry (ICP-MS). (E) Scatter dot plot with bar at mean ratio of Cu detected by ICP-MS to protein ± s.e.m. of MEK1, as isolated, or MEK1 positive fractions 15 and 16 from Cu-CCS and MEK1 mixed 1:1. n=2 independent experiments. Results were compared using a unpaired, two-tailed Student’s t-test. **, P=0.0081. (F) Scatter dot plot with bar at mean luminescence units ± s.e.m. from ELISA detection of recombinant phosphorylated (P)-ERK2^K54R^ from MEK1^WT^, MEK1^H188A^(H188A), MEK1^H188F^(H188F), MEK1^H188M^(H188M), MEK1^C207A^(C207A), or MEK1^C207S^(C207S) incubated with increasing concentrations of Cu-CCS. n=4 technical replicates or MEK1^H188M^ n=2 technical replicates. Results were compared using a two-way ANOVA followed by a Tukey’s multi-comparisons test. *, P=0.0147; ****, P<0.0001.

We next interrogated the relationship between the Cu chaperones and MEK1 kinase activity and Cu binding *in vitro*. A selective 3-fold enhancement in MEK1 kinase activity was observed when increasing concentrations of Cu loaded CCS but not ATOX1 were added to MEK1 kinase assays (Figure 4C), indicating that CCS is directly involved in Cu-stimulated MEK1/2 phosphorylation of ERK1/2(Brady et al., 2014). Based on the observation that CCS, which binds Cu(I), increased MEK1 kinase activity, we evaluated whether this increase in activity was driven by Cu transfer. Stoichiometric addition of Cu-CCS to MEK1 and subsequent separation by SEC to capture a predominantly MEK1 fraction for analysis by ICP-MS resulted in an average increase of 1.2 Cu atoms bound per MEK1 protomer (Figures 4D and 4E). Finally, we sought to connect CCS-mediated Cu transfer to increased MEK1 kinase activity to the coordinating ligands revealed in the Cu-MEK1 X-ray crystal structure. As expected, Cu-CCS failed to increase the kinase activity of either MEK^H188A^, MEK1^H188F^, or MEK^C207A^ when compared to wild-type (Figure 4F), suggesting that the Cu(II) coordinating ligands depicted in the X-ray crystal structure (Figure 1) are necessary for CCS delivery of Cu(I) to MEK1, which activates the kinase. Further, Cu-CCS stimulation of MEK1 mutants, in which H188 or C207 were substituted with weakly Cu coordinating amino acids(Rubino and Franz, 2012), was equivalent to wild-type MEK1 basal kinase activity levels (Figure 4F). These data infer an intricate link between Cu(I) coordination and basal level activity that can be achieved when the molar ratio of Cu(I) bound MEK1 is selectively increased within the MEK1 active site. Taken together, these data support a scenario in which the cytosolic Cu chaperone CCS functions as a direct Cu delivery mechanism to MEK1 via a transient association that in turn facilitates increased kinase activity.

### Cu integration into MAPK signaling requires CCS Cu binding

To directly address an association between CCS and MAPK signaling in mammalian cells, BirA proximity-dependent biotin identification (BioID) reagents were generated to capture the weak or transient interaction between ATOX1 or CCS with MEK1 in living cells. Specifically, myc-epitope tagged mutant (R118G) BirA biotin ligase, which conjugates biotin to proteins within the immediate vicinity (∼10 nm)(Kim et al., 2016), was fused in frame to the N-terminus of ATOX1 or CCS and stably expressed in HEK-HT cells (Figure 5A). As expected, exogenous biotin treatment of HEK-HT cells expressing myc-BirA-ATOX1 resulted in the streptavidin recovery of biotinylated ATP7A(Kim et al., 2008; Robinson and Winge, 2010) (Figure 5A), which facilitates the transport of Cu into the lumen of the trans-Golgi network where Cu loading of Cu-dependent enzymes occurs, while myc-BirA-CCS biotinylated SOD1(Culotta et al., 1997; Kim et al., 2008; Robinson and Winge, 2010) (Figure 5A), which utilizes Cu as a cofactor for catalyzing the disproportionation of superoxide to hydrogen peroxide and dioxygen. Endogenous MEK1 was only biotinylated in presence of biotin and myc-BirA-CCS, indicating that CCS is proximal to MEK1/2 and may be responsible for Cu-stimulated activation of MAPK signaling.

**Figure 5.**
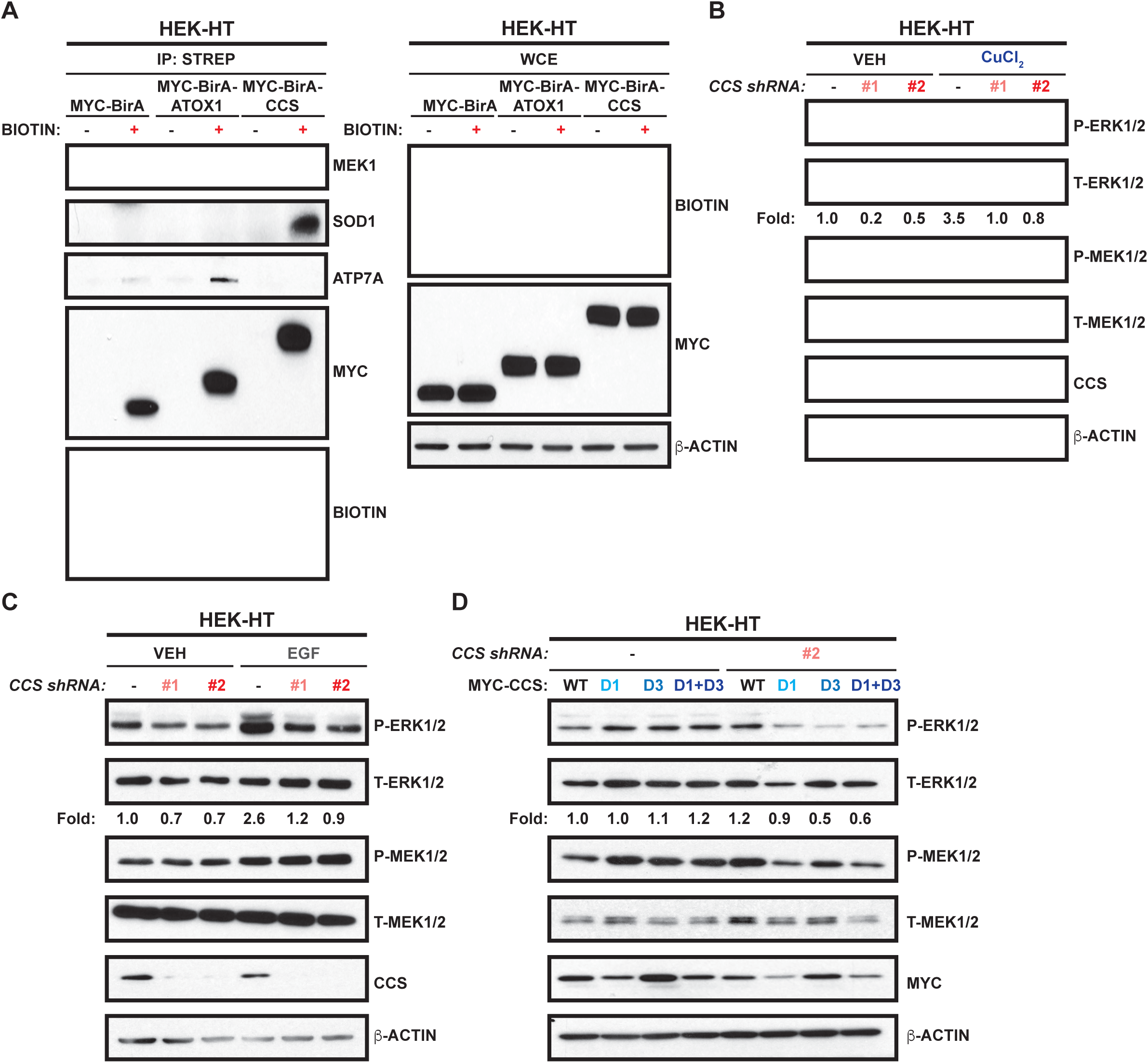
CCS is the necessary Cu chaperone for MEK1. (A) Immunoblot detection of biotinylated MEK1, ATP7A, SOD1, MYC-BirA, MYC-BirA-ATOX1, or MYC-BirA-CCS from strepavadin pulldowns and whole cell extracts from HEK-HT cells stably expressing *MYC-BirA, MYC-BirA-ATOX1*, or *MYC-BirA-CCS* treated with media with (+) or without (-) biotin for 48 hours. n=2 biologically independent experiments. (B and C) Immunoblot detection of phosphorylated (P)-ERK1/2, total (T)-ERK1/2, P-MEK1/2, T-MEK1/2, CCS, or β-ACTIN from HEK-HT cells stably expressing doxycycline inducible *shRNA* against control (-) or *CCS* (*#1 or #2*) treated with doxycycline for 24 hours followed by (B), vehicle (VEH) or 1µM CuCl_2_ for 20 minutes or (C), VEH or 0.1ng/mL EGF for 15 minutes. Quantification: Fold change P-ERK1/2/T-ERK1/2 normalized to control, VEH. n=3 biologically independent experiments. (D) Immunoblot detection of P-ERK1/2, T-ERK1/2, P-MEK1/2, T-MEK1/2, MYC, or β-ACTIN from HEK-HT cells stably expressing doxycycline inducible *shRNA* against control (-) or *CCS* (*#2*) reconstituted with *MYC-CCS*^*WT*^(WT), *MYC-CCS Domain 1 Mutant* (D1), *MYC-CCS Domain 3 Mutant* (D3), or *MYC-CCS Domain 1+3 Mutant* (D1+D3) treated with doxycycline for 24 hours. Quantification: Fold change P-ERK1/2/T-ERK1/2 normalized to control, VEH. n=3 biologically independent experiments.

To investigate whether CCS Cu binding is necessary for endogenous MAPK signaling, *CCS* was knocked down with two independent doxycycline-inducible shRNAs in HEK-HT cells. Knockdown of *CCS* decreased basal and either CuCl_2_ or EGF stimulated activation of P-ERK1/2 (Figures 5B and 5C). CCS-mediated metalation of SOD1 involves three distinct domains of CCS that are each required for SOD1 activity. Domain 1 (D1) facilitates Cu acquisition, domain 2 (D2) mimics SOD1 to promote protein-protein interactions, and domain 3 (D3) may be involved in D1 to D3 Cu transfer while also generating the disulfide bond necessary for SOD1 activation(Boyd et al., 2019; Fetherolf et al., 2017; Robinson and Winge, 2010). To definitively test the contribution of CCS D1 and D3 to MAPK signaling, HEK-HT stably expressing non-targeting control or doxycycline-inducible *CCS* shRNA in the presence of exogenous wild-type or mutant CCS at the Cu binding interface of D1, D3, or both were generated (Figure 5D). Based on its inability to acquire Cu regardless of the source, mutation of CCS D1 resulted in reduced P-ERK1/2 (Figure 5D). Surprisingly, D3 of CCS was also required for basal P-ERK1/2 levels (Figure 5D), suggesting that a similar mechanism by which CCS activates SOD1 may be utilized to facilitate Cu-dependent CCS-mediated activation of MEK1/2(Boyd et al., 2019; Fetherolf et al., 2017) and is supported by the combined D1/D3 mutant not exhibiting an exacerbated phenotype. Taken together, these results indicate that CCS is an integral component of MEK1/2 responsiveness to both Cu and growth factors.

### Targeting CCS with the small molecule inhibitor DCAC50 reduces MEK1/2 activity

A recent study described the development and characterization of a small molecule inhibitor, DCAC50, targeting the Cu chaperones ATOX1 and CCS(Wang et al., 2015). Mechanistically, DCAC50 binds ATOX1 and CCS with K_D_ of 6.8±1.7µM and 8.2±2.7µM, respectively, and prevents Cu transfer between the chaperones and their downstream interacting target proteins(Wang et al., 2015). Thus, to investigate whether acute inhibition of CCS would decrease MAPK pathway activation in an analogous fashion to genetic deletion of *CCS* (Figures 5B and 5C), HEK-HT cells were treated with increasing concentrations of DCAC50. Similar to loss of *CCS*, P-ERK1/2 was reduced in response to DCAC50 treatment in a dose-dependent fashion (Figure 6A). Further, the kinetics of MAPK pathway activation by CuCl_2_ or EGF were dampened and delayed in the presence of DCAC50 (Figures 6B and 5C). While knockout of *ATOX1* in human BRAF mutation positive melanoma cells also decreased MAPK signaling and DCAC50 targets both CCS and ATOX1, exogenous CuCl_2_ was sufficient to rescue P-ERK1/2(Kim et al., 2019), indicating that ATOX1 impacts MAPK pathway activation indirectly of facilitating Cu binding of MEK1/2. These results support our findings that CCS is directly required for MAPK signaling by contributing to MEK1/2 Cu binding and suggest another avenue to pharmacologically target aberrant MAPK signaling required for malignant transformation (Figure 6D).

**Figure 6.**
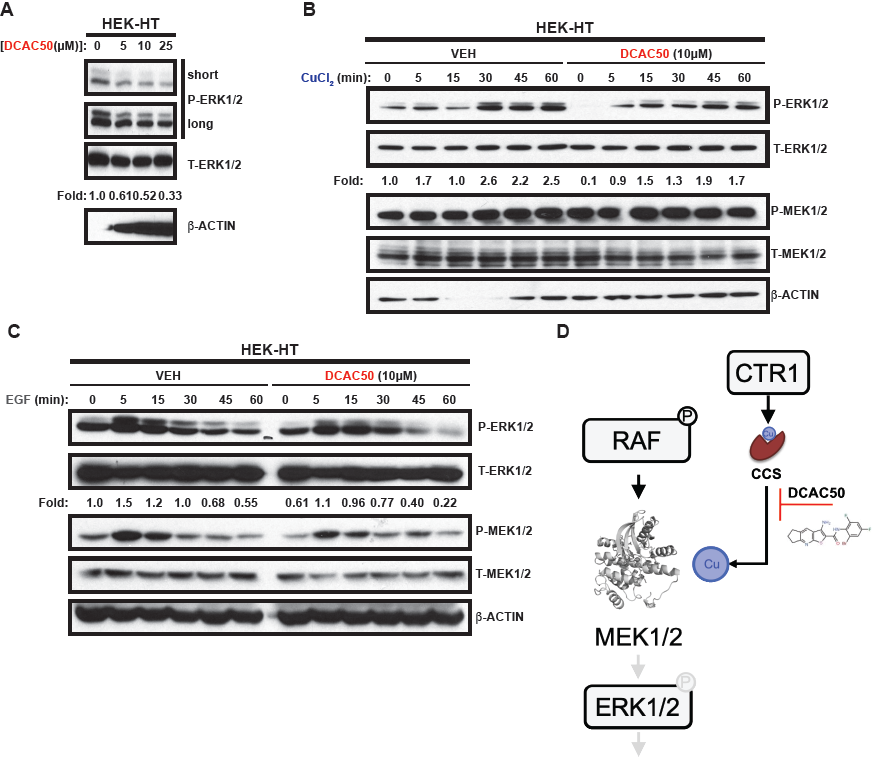
Treatment with small molecular inhibitor of CCS blunts MAPK pathway activation at the level of MEK1/2. (A) Immunoblot detection of phosphorylated (P)-ERK1/2, total (T)-ERK1/2, or β-ACTIN from HEK-HT cells treated with vehicle (0) or increasing concentrations of DCAC50 for 24 hours. Quantification: Fold change P-ERK1/2/T-ERK1/2 normalized to control, VEH. n=3 biologically independent experiments. (B and C) Immunoblot detection of P-ERK1/2, T-ERK1/2, P-MEK1/2, T-MEK1/2, or β-ACTIN from HEK-HT cells treated with vehicle (-) or 10µM DCAC50 for 24 hours followed stimulation with (B), 1µM CuCl_2_ or (C), 0.1ng/mL EGF for indicated time. Quantification: Fold change P-ERK1/2/T-ERK1/2 normalized to control, VEH. n=3 biologically independent experiments. (D) Model of CCS-mediated Cu activation of MEK1/2.

## Discussion

Our previous discovery that Cu selectively regulates the canonical MAPK pathway at the level of the MEK1/2 kinases, in both *Drosophila* and mammalian cell settings suggests that there is an evolutionarily conserved pressure for this integration(Turski et al., 2012). Further, it established a critical mechanistic function for Cu as an intracellular mediator of signaling, which was previously reserved for redox-inactive metals like Zn^2+^, K^+,^ Na^+^, and Ca^2+^ that have well-appreciated roles in cell signaling(Chang, 2015). However, the structural and molecular level basis for Cu-dependent MEK1/2 kinase activity remained to be interrogated. We provide here the first X-ray crystal structure of a Cu bound mammalian kinase and demonstrate that MEK1 coordinates Cu at critical amino acids within the MEK1 active site (Figure 1). Large rearrangements between the apo- and Cu-MEK1 structures were absent (Table 1) and preclude a definitive mechanistic explanation of Cu-stimulated kinase activity (Figure 4). Nevertheless, two of the chemical ligands (H188 and C207) responsible for Cu(II) coordination (Figure 1) and Cu(I) affinity (Table 4) were essential for downstream phosphorylation of ERK1/2 and solidified their importance (Figure 3). While numerous X-ray crystal structures of MEK1 have been previously resolved, none contain Cu despite sub femtomolar affinity for full length MEK1(Turski et al., 2012). This suggests that purified MEK1 is devoid of significant levels of Cu, which is supported by the low stoichiometry of Cu in the isolated protein (Figure 4E)(Turski et al., 2012). This result could also suggest that crystallizable MEK1 fragments may have reduced Cu affinity and/or the crystallization conditions failed to capture the Cu bound state. Metalloregulation can be achieved through kinetically labile but thermodynamically stable sites(Foster et al., 2014) and the alternate 20% occupancy within the crystal structures may be a result of different Cu bound oxidation states and/or movement within the active site (Figure S1). Interestingly, the chemical ligands revealed in the X-ray crystal structure (Figure 1) do not entirely overlap with a stretch of residues (M187, H188, M230, H239) within the MEK1 kinase domain previously identified by targeted mutagenesis and a metal-catalyzed oxidation reaction followed by mass spectrometry to be critical for Cu binding and kinase activity(Brady et al., 2014). While M187, M230, and H239 do not participate in Cu ligation in the X-ray crystal structure, the region adjacent to or containing the residues M230 and H239 are moreover unresolved in the X-ray crystal structure, thus obscuring their roles in Cu binding and/or activation. Both MEK1 and MEK2: *i*) bind Cu, *ii*) kinase activity can be enhanced with Cu and silver, which is isoelectric to Cu(I), and *iii*) are conserved at amino acids identified in the X-ray crystal structure (H188, C207, D208) and previously (M187, H188, M230, H239) (Brady et al., 2014). In contrast, MEK5, which is highly similar to MEK1/2,: *i*) fails to bind Cu, *ii*) kinase activity is not responsive to Cu or Cu chelator, and *iii*) lacks conservation of residues M230 and H239 but retains the residues that mediate Cu(II) ligation in the X-ray crystal structure amino acids(Brady et al., 2014). Importantly, introduction of analogous mutations in MEK5 (H281 and M323), which resulted in the generation of a Cu binding mutant-equivalent MEK5, did not alter MEK5 kinase activity towards the common kinase substrate MBP or the MEK5-specific substrate ERK5 when compared to wild-type MEK5(Brady et al., 2014). Thus, we hypothesize that MEK1 interacts with Cu at two distinct sites of which the first (involving M230 and H239) is an intermediate, “entry site” that is essential for handing off Cu to the active site (involving H188 and C207) and thus, both are critical for Cu(I)-stimulated kinase activation (Figure 1). Finally, the chemical environment of MEK1 Cu(II) coordination was corroborated in extensive spectroscopy studies, indicating the X-ray crystal structure is representative of an active Cu-MEK1 species (Figure 2).

Intriguingly, spectroscopy studies revealed an interesting MEK1 Cu binding mode suggestive of a reductive mechanism that generates Cu(I) *in situ* (Figure 2). When Cu(II) was provided to MEK1 in an aerobic environment, MEK1 bound Cu(II) readily and reduced it to Cu(I) immediately. These findings suggest that MEK1 has the ability to change the oxidation state of Cu bound and this is presumably related to the oxidation state of MEK1 itself. One possible mechanism to change the oxidation state of bound Cu is through a cluster of cysteine side-chains, like C121, C142 and C207, whose oxidation could provide the electrons necessary for Cu(II) reduction. This process has been fully characterized in the formation of the mixed-valence di-Cu center in the CuA site of subunit 2 of cytochrome oxidase(Chacón and Blackburn, 2012) and in the formation of the similar site in purple CuA azurin(Chakraborty et al., 2015), where disulfide formation in one-third of the population of subunits provides the electrons necessary for formation of the Cu(I) atoms that bind to the other two-thirds of the protein. The process leads to a characteristic sub-stoichiometric Cu to protein ratio (0.67 CuA centers per subunit) since molecules where the thiolate ligands have oxidized to the disulfide state cannot form the binuclear mixed-valence site. It is probable that chemistry similar to this is occurring in MEK1, which would explain why the Cu to protein ratios in wild-type are sub stoichiometric, as well as the resulting Cu(I)-bound state. However, the observed ratio (0.5 Cu/P) deviates from the expected 0.67 leaving open other possibilities. One scenario that would lead to 0.5 Cu/P ratio would be disulfide formation between two cysteine residues one from each monomer to produce a disulfide-bridged dimer where a single Cu(I) could bind at the dimer interface. This situation has been found in other Cu(I) trafficking and/or sensing kinases such as CusS(Affandi et al., 2016). The EXAFS simulations with 2 Cu(I)-S interactions (Table 3) can accommodate such a model, giving equivalent fit quality in some but not all of the samples. We also note that a coordination site composed of the thioether S atoms from M187 residues, which were previously identified to be oxidized in a Cu-dependent fashion following metal catalyzed oxidation reactions, of each monomer would be consistent with our data, and that Met-bridged dimers have been characterized by crystallography in other Cu systems. More definitive studies to interrogate the existence of Cu-meditated MEK1 disulfide and its contribution to MEK1 activation are warranted.

In support of the hypothesis that kinases within complex protein modules may directly respond to and/or sense metals like Cu, we found that short exposure to CuCl_2_ was sufficient to robustly increase ERK1/2 phosphorylation (Figures 3 and 5). This observation indicates that MEK1/2 are not fully Cu bound in cells and are thus primed to rapidly respond to fluctuations in intracellular Cu levels. The cellular contexts in which a labile Cu pools is liberated are under active investigation. Two recent reports provide evidence of this phenomena in which depletion of the intracellular glutathione pools(Chung et al., 2019) or amino acid deprivation(Tsang et al., 2020) were sufficient to elevate Cu(I) levels. Further, both basal and growth factor stimulated MAPK signaling were diminished when point mutants of the Cu coordinating ligands were introduced, which suggests that Cu is a hardwired component of the pathway that is necessary for the other inputs.

To balance cellular toxicity from free Cu ions with metalation of cuproenzymes, evolutionarily conserved Cu chaperones promote Cu distribution. Here, we show that the Cu chaperone for superoxide dismutase CCS transiently binds to and facilities Cu loading into MEK1 and in turn increases MEK1/2 kinase activity (Figure 4). In agreement with a whole genome RNAi screen in *Drosophila* S2 cells aimed at identifying modulators of MAPK signaling(Friedman and Perrimon, 2006b), knockdown of *CCS* reduced ERK1/2 phosphorylation (Figure 5). While the X-ray crystal structure exhibits a square planar arrangement of the Cu atom, indicative of Cu(II) binding (Figure 1), the biochemical studies support a biologically relevant, CCS-mediated Cu(I) activation of MEK1 that requires H188 and C207 (Figure 5). Similar to SOD1(Fetherolf et al., 2017), which is also loaded with Cu(I) by CCS, MEK1 may bind to both CCS and Cu at an intermediate “entry site” involving M187, M230, and H239 that provides specificity and is necessary for eventual Cu coordination within the kinase active site at H188, C207, and D208 (Figure 1). Specifically, an intricate molecular mechanism requiring three distinct domains of CCS is required for SOD1 maturation and activation(Boyd et al., 2019; Fetherolf et al., 2017; Robinson and Winge, 2010). Interestingly, D3 of CCS, which helps with Cu transfer into the SOD1 active site and generates a disulfide bond in SOD1(Boyd et al., 2019; Fetherolf et al., 2017; Robinson and Winge, 2010), was also necessary for robust MEK1/2 activation (Figure 5). This finding suggests that the aforementioned Cu-mediated MEK1 disulfide formation may be installed by CCS in mammalian cells and mechanistically underlie MEK1 activation by Cu. Interestingly, *in vitro* experiments adding CuCl_2_ to purified MEK1 protein resulted in inhibition, not activation, of the enzyme (data not shown), suggesting that Cu loading by CCS is required for activation, potentially to avoid Cu binding to other sites that might inhibit MEK1 activity or to facilitate complete maturation of the Cu bound state. This is consistent with the observation that the MEK1 crystals soaked with CuCl_2_ in the absence of CCS showed weak anomalous signals near residues Met 187 in chain A and Cys121 in chain B (*see* crystallography methods), which could represent additional Cu binding sites correlated with MEK1 inhibition. This result further supports the importance of CCS in controlling MEK1 activation via Cu.

Finally, pharmacologic targeting of CCS with the small molecule inhibitor DCAC50(Wang et al., 2015) blunted both basal and stimulated activation of MEK1/2 (Figure 6). The vital importance of intricate mechanisms to dynamically regulate MAPK signaling output is underscored by its dysregulation in approximately 85% of human cancers(Caunt et al., 2015; Holderfield et al., 2014). While targeting the protein kinase catalytic activity of MEK1/2 is approved in the setting of BRAF^V600E^ metastatic melanoma(Caunt et al., 2015; Flaherty et al., 2012) and is under investigation in several other cancers, resistance mechanisms limited clinical durability. Therefore, our structural and molecular studies provide a framework for the developmental of MEK1/2 inhibitors that exploit its Cu dependence by targeting the binding interface or intracellular delivery components.

## Acknowledgements

We thank C.M. Counter (Duke University), J. Chen (Emory University), and P. Hart (University of Texas Health Sciences Center San Antonio) for reagents, L.A. Brady, L. Busino, M.L. Matthews, and E.S. Witze for technical support, discussions, and/or review of the manuscript, and D. Sneddon for administrative support. This work was supported by NIH grants GM124749 (D.C.B.), CA243294 (T.T), GM123725 (N.J.B), and GM136239 (N.J.B.), Swedish Research Council Grant 2015-03881 (P.W.S.), Knut and Alice Wallenberg Foundation Scholar Grant (P.W.S.), and Pew Scholars Program in Biomedical Science Award #50359 (D.C.B.). Use of the Advanced Photon Source was supported by the U. S. Department of Energy, Office of Science, Office of Basic Energy Sciences, under Contract No. DE-AC02-06CH11357. Data was collected on beamline 24-ID-C at the Advanced Photon Source, Argonne National Laboratory. Use of the Stanford Synchrotron Radiation Lightsource (SSRL), SLAC National Accelerator Laboratory, is supported by the U.S. Department of Energy, Office of Science, Office of Basic Energy Sciences under Contract No. DE-AC02-76SF00515. The SSRL Structural Molecular Biology Program is supported by the DOE Office of Biological and Environmental Research, and by NIH, NIGMS (including P41 GM103393).

## Author Contributions

G.J.B. and Y.J.K. contributed equally to this work. M.G. contributed to the study design, generated plasmids and recombinant proteins, performed X-ray crystallography, *in vitro* kinase assays, size exclusion chromatography, analyzed data, prepared figures, and contributed to writing manuscript. G.J.B. generated plasmids and cell lines, performed Western blot analysis, analyzed data, and contributed to figure preparation. Y.J.K. generated cell lines, performed Western blot analysis, analyzed data, and contributed to figure preparation. K.B.A. performed EPR, XAS, and EXAFS, analyzed data, and contributed to figure preparation. S.B. generated recombinant proteins, performed Cu(I) BCA affinity, electrophoretic mobility shift assays, and size exclusion chromatography, analyzed data, and contributed to figure preparation. M.M.D. and S.V. generated recombinant protein, performed surface plasmon resonance assays, analyzed data, and contributed to figure preparation. T.T. performed co-immunoprecipitations and Western blot analysis, analyzed data, and contributed to figure preparation. N.A.S. generated plasmids and recombinant proteins, assisted X-ray crystallography, *in vitro* kinase assays, size exclusion chromatography, and analyzed data. M.L.M. provided expertise in metal coordination chemistry and contributed to writing manuscript. G.M.B. provided expertise in X-ray crystal structure modeling and small molecule inhibitor synthesis, and contributed to writing manuscript. P.W.S. contributed to study design, provided expertise in biochemical and biophysical assays, facilitated data analysis, and contributed to writing manuscript. D.D.W. contributed to study design, provided expertise in biochemical and biophysical assays, facilitated data analysis, and contributed to writing manuscript. N.J.B. contributed to study design, provided expertise in biochemical and biophysical assays, facilitated data analysis, and contributed to writing manuscript. R.M. contributed to study design, provided expertise in X-ray crystallography, biochemical assays, facilitated data analysis, and contributed to writing manuscript. D.C.B conceived of the project, contributed to the study design, generated cell lines, plasmids, and recombinant proteins, analyzed data, prepared figures, and wrote the manuscript. All authors read and provided feedback on manuscript and figures.

## STAR Methods

### Cell lines

HEK-HT cells were previously described(Hahn et al., 1999) and provided by C.M. Counter (Duke University). HEK-HT cells were maintained in Dulbecco’s Modified Eagle Media (DMEM, Gibco) supplemented with 10% v/v fetal bovine serum (FBS, GE Lifesciences) and 1% penicillin-streptomycin (P/S, Gibco). HEK-HT cells were stably infected with lentiviruses derived from the pSMARTvector inducible lentiviral shRNA plasmids (Dharmacon, *see* plasmids below) or retroviruses derived from the pWZL retroviral plasmid (*see* plasmids below) described below using established methods.

### Plasmids

pGEX4T3-HA-ERK2^K54R^ (Addgene plasmid #53170)(Brady et al., 2014), pET3A-T7-HIS-ATOX1(Horvath et al., 2019), pWZLblasti-HA-MEK1^WT^ (Addgene plasmid #53161)(Brady et al., 2014), and pET15B-HIS-PARVALBUMIN(Werner et al., 2018) were previously described. pET28A-6XHIS-TEV-MEK1^(45-393/Δ264-307/SVQSDI)^ was created by PCR subcloning with primers designed against *MEK1* cDNA sequence corresponding to the amino acids 45-393 and subsequently replacing sequence corresponding to the amino acids 264-307 with a 6-residue sequence SVQSDI from a *MAP2K1* cDNA clone (Dharmacon # MHS 6278-211690391) for the pET28A plasmid containing an N-terminal, TEV protease cleavable 6XHis tag. pET28A-6XHIS-TEV-MEK1^WT^ was created by PCR subcloning MEK1^WT^ from a *MAP2K1* cDNA clone (Dharmacon # MHS 6278-211690391) with primers for the pET28A plasmid containing an N-terminal, TEV protease cleavable 6XHis tag. pET28A-6XHIS-TEV-MEK1^H188A^, -MEK1^H188F^, – MEK1^H188M^, -MEK1^C207A^, and -MEK1^C207S^ were created by introducing mutations corresponding to the indicated amino acid changes by site-directed mutagenesis into MEK1^WT^ from the pET28A-6XHIS-TEV-MEK1^WT^ plasmid. pWZLblasti-HA-MEK1^H188A^ and -MEK1^C207A^ were created by introducing mutations corresponding to the indicated amino acid changes by site-directed mutagenesis into MEK1^WT^ from the pWZLblasti-HA-MEK1^WT^ plasmid. pETDUET-6XHIS-TEV-ATOX1 and pBABEpuro-MYC-BioID2-ATOX1 were created by PCR subcloning ATOX1 from pCDNA-ATOX1(Wang et al., 2015) with primers for the pETDUET plasmid containing an N-terminal, TEV protease cleavable 6XHIS-tag or the pBABEpuro-MYC-BioID2 (Addgene #80900)(Kim et al., 2016) plasmid containing an N-terminal myc tag in frame with mutant BirA (R118G). pETDUET-6XHIS-TEV-CCS, pBABEpuro-MYC-BioID2-CCS, and pWZLblasti-MYC-CCS^WT^ were created by PCR subcloning CCS from pCDNA-CCS(Wang et al., 2015) with primers for the pETDUET plasmid containing an N-terminal, TEV protease cleavable 6XHIS-tag, for the pBABEpuro-MYC-BioID2 plasmid containing an N-terminal MYC-tag in frame with mutant BirA (R118G), or with primers including an N-terminal MYC-tag for the pWZLblasti plasmid. pKA8H-CCS was created by PCR subcloning CCS with primers for the pKA8H plasmid containing an N-terminal, TEV protease cleavable 8XHIS-tag. pWZLblasti-MYC-CCS^D1(C22A/C25A)^, -CCS^D3(C244A/C246A)^, and -CCS^D1+D3(C22A/C25A/C244A/C246A)^ were created by introducing mutations corresponding to the indicated amino acid changes by site-directed mutagenesis into CCS^WT^ from the pWZLblasti-MYC-CCS^WT^ plasmid. pSMARTvector inducible lentiviral shRNA plasmids were purchased from Dharmacon to express: nontargeted control, human CCS target sequence #1 5’-GGGCAGGCAAAGTCCCTCT and human CCS target sequence #2 5’-AGTGGAAAGTGCTCGCCCT. GFP and shRNA expression was induced by adding 2 µg/ml doxycycline hydrochloride (Fisher Scientific, AAJ67043AE) to DMEM supplemented with 10% FBS and 1% P/S.

### Protein Expression and Purification

#### Proteins used X-ray crystallography

##### 6XHIS-TEV-MEK1^(45-393/Δ264-307/SVQSDI)^

The MEK1 construct spanning residues 45-393 and with replacement of the flexible linker region 264-307 with a 6-residue sequence SVQSDI (MEK1^45-393/ Δ264-307/SVQSDI^) was expressed in BL21 (DE3) RIL cells in Terrific Broth (TB, Dot Scientific) at 37 °C until reaching an OD600 of 0.800 and then induced with 1 mM isopropyl β-D-1-thiogalactopyranoside (IPTG) overnight at 17 °C. The cells were then spun down the next day and lysed in lysis buffer 1 (25 mM Tris pH 7.5, 500 mM NaCl, and 5 mM BME) treated with 1 mM PMSF and DNAseI. The lysate was spun down at 19,000 r.p.m. for 40 minutes and the supernatant was added to 10 mL of HisPur Ni-NTA resin (ThermoFisher Scientific, # 88223) previously washed with lysis buffer 1 and left to incubate at 4 °C for 1 hr. The supernatant was eluted via gravity column and the resin was then washed with 1 L of lysis buffer 1 treated with 20 mM imidazole. The protein was then eluted with lysis buffer 1 supplemented with 300 mM imidazole and the elution fractions were pooled together and treated with TEV protease overnight while dialyzing into dialysis buffer 1 (25 mM Tris pH 7.5, 20 mM NaCl, 5 mM BME). The eluent is then applied to a HiTrap SP HP cation exchange 5mL column and eluted over a gradient of 20 column volumes ranging from 0% to 100% of Buffer B (25 mM Tris pH 7.5, 1 M NaCl, and 5 mM BME). The protein elutes between 10% and 40% Buffer B. Peak fractions were run on an SDS-PAGE gel, pooled, concentrated, and applied to 10 mL of fresh Ni-NTA resin washed with reverse nickel buffer 1 (25 mM Tris pH 7.5, 250 mM NaCl, 20 mM imidazole, and 5 mM BME). The protein was allowed to pass over the nickel resin 3 times, and then washed extensively with reverse nickel buffer 1 until no more protein was washed off the resin as monitored by Bradford Reagent (Sigma-Aldrich, # B6916). The protein was then concentrated and applied to a Superdex S200 10/300 gel filtration column in a final buffer of 20 mM HEPES pH 7.0, 150 mM NaCl, 5 mM BME. Protein was concentrated to 10 mg/mL (∼285 µM) and immediately used for crystallization.

#### Proteins used for in vitro kinase assays, XAS, or EXAFS

##### 6XHIS-MEK^WT^, -MEK1^H188A^, -MEK1^H188F^, -MEK1^H188M^, -MEK1^C207A^, and -MEK1^C207S^

6XHIS-MEK^WT^ and mutants (H188A, H188F, H188M, C207A, and C207S) were expressed in BL21 (DE3) RIL cells in TB at 37 °C until reaching an OD600 of 0.700 and then induced with 1 mM IPTG overnight at 17 °C. The cells were then spun down the next day and lysed in lysis buffer 1 (25 mM Tris pH 7.5, 500 mM NaCl, and 5 mM BME) treated with 1 mM PMSF and DNAseI. The lysate was then spun down at 19,000 r.p.m. for 40 minutes and the supernatant was added to 10 mL of HisPur Ni-NTA resin previously washed with lysis buffer 1 and left to incubate at 4 °C for 1 hr. The supernatant was eluted via gravity column and the resin was then washed with 1 L of lysis buffer 1 treated with 20 mM imidazole. The protein was then eluted with lysis buffer 1 supplemented with 300 mM imidazole and the elution fractions were pooled together and treated with TEV protease overnight while dialyzing into dialysis buffer 1 (25 mM Tris pH 7.5, 20 mM NaCl, 5 mM BME). The eluent is then applied to a HiTrap Q HP anion exchange 5mL column and eluted over a gradient of 20 column volumes ranging from 0% to 100% of Buffer B (25 mM Tris pH 7.5, 1 M NaCl, and 5 mM BME). The protein elutes between 15% and 30% Buffer B. Peak fractions were run on an SDS-PAGE gel, pooled, concentrated, and applied to 10 mL of fresh Ni-NTA resin washed with reverse nickel buffer 1 (25 mM Tris pH 7.5, 250 mM NaCl, 20 mM imidazole, and 5 mM BME). The protein was allowed to pass over the nickel resin 3 times, and then washed extensively with reverse nickel buffer 1 until no more protein was washed off the resin as monitored by Bradford Reagent. The protein was concentrated and run on a Superdex S200 10/300 gel filtration column in a final buffer of 20 mM HEPES pH 7.0, 150 mM NaCl, 5 mM BME. Protein was concentrated to 2-5 mg/mL (80-200 µM), flash frozen in liquid nitrogen, and stored in −80 °C freezer for future use.

##### GST-ERK2^K54R^

GST-ERK2^K54R^ was expressed in BL21 (DE3) RIL cells in TB at 37 °C until reaching an OD600 of 0.700 and then induced with 1 mM IPTG overnight at 17 °C. The cells were then spun down the next day and lysed in lysis buffer 1 (25 mM Tris pH 7.5, 500 mM NaCl, and 5 mM BME) treated with 1 mM PMSF and DNAseI. The lysate was then spun down at 19,000 r.p.m. for 40 minutes and the supernatant was added to 10 mL of Glutathione Agarose Resin (Pierce, 16102BID) previously washed with lysis buffer 1 and left to incubate at 4 °C for 2 hr. The supernatant was eluted via gravity column and the resin was then washed with 1 L of lysis buffer 1. The protein was then eluted with lysis buffer 1 supplemented with 20 mM glutathione (Sigma Aldrich # G4521) and the elution fractions were pooled together and dialyzed into dialysis buffer 1 (25 mM Tris pH 7.5, 150 mM NaCl, 5 mM BME). The protein was then concentrated and applied to a Superdex S200 10/300 gel filtration column in a final buffer of 25 mM Tris pH 7.5, 150 mM NaCl, 5 mM BME. Protein was concentrated to 2.4 mg/mL (35 µM), flash frozen in liquid nitrogen, and stored in −80 °C freezer for future use.

##### 6XHIS-ATOX1

6XHIS-ATOX1 was expressed in BL21 (DE3) RIL cells in LB (Millipore) at 37 °C until reaching an OD600 of 0.700 and then induced with 1 mM IPTG overnight at 17 °C. The cells were then spun down the next day and lysed in lysis buffer 1 (25 mM Tris pH 7.5, 500 mM NaCl, and 5 mM BME) treated with 1 mM PMSF and DNAseI. The lysate was then spun down at 19,000 r.p.m. for 40 minutes and the supernatant was added to 10 mL of HisPur Ni-NTA resin previously washed with lysis buffer 1 and left to incubate at 4 °C for 1 hr. The supernatant was eluted via gravity column and the resin was then washed with 1 L of lysis buffer 1 treated with 20 mM imidazole. The protein was then eluted with lysis buffer 1 supplemented with 300 mM imidazole and the elution fractions were pooled together and treated with TEV protease overnight while dialyzing into dialysis buffer 1 (25 mM Tris pH 7.5, 20 mM NaCl, 5 mM BME). The eluent is then applied to a HiTrap Q HP anion exchange 5mL column and eluted over a gradient of 20 column volumes ranging from 0% to 100% of Buffer B (25 mM Tris pH 7.5, 1 M NaCl, and 5 mM BME). Peak fractions were run on an SDS-PAGE gel, pooled, concentrated, and applied to 10 mL of fresh Ni-NTA resin washed with reverse nickel buffer 1 (25 mM Tris pH 7.5, 250 mM NaCl, 20 mM imidazole, and 5 mM BME). The protein was allowed to pass over the nickel resin 3 times, and then washed extensively with reverse nickel buffer 1 until no more protein was washed off the resin as monitored by Bradford Reagent. The protein was then concentrated and added to a Superdex S200 10/300 gel filtration column in a final buffer of 20 mM HEPES pH 7.0, 150 mM NaCl, 5 mM BME. Protein was concentrated to ∼6 mg/mL (∼650 µM) and dialyzed into dialysis buffer 2 (20 mM HEPES pH 7.0, 150 mM NaCl, 3.25 mM DTT) overnight. The next morning 1M CuCl_2_ was added to 650 µM of ATOX1 to a final concentration of 585 µM of CuCl_2_ and mixed until dissolved. The protein loaded with Cu was then flash frozen in liquid nitrogen and stored in −80 °C freezer for future use.

##### 6XHIS-CCS

6XHIS-CCS was expressed in BL21 (DE3) RIL cells in LB at 37 °C until reaching an OD600 of 0.700 and then induced with 1 mM IPTG overnight at 17 °C. The cells were then spun down the next day and lysed in lysis buffer 1 (25 mM Tris pH 7.5, 500 mM NaCl, and 5 mM BME) treated with 1 mM PMSF and DNAseI. The lysate was then spun down at 19,000 r.p.m. for 40 minutes and the supernatant was added to 10 mL of HisPur Ni-NTA resin previously washed with lysis buffer 1 and left to incubate at 4 °C for 1 hr. The supernatant was eluted via gravity column and the resin was then washed with 1 L of lysis buffer 1 treated with 20 mM imidazole. The protein was then eluted with lysis buffer 1 supplemented with 300 mM imidazole and the elution fractions were pooled together and treated with TEV protease overnight while dialyzing into dialysis buffer 1 (25 mM Tris pH 7.5, 20 mM NaCl, 5 mM BME). The eluent is then applied to a HiTrap Q HP anion exchange 5mL column and eluted over a gradient of 20 column volumes ranging from 0% to 100% of Buffer B (25 mM Tris pH 7.5, 1 M NaCl, and 5 mM BME). Peak fractions were run on an SDS-PAGE gel, pooled, concentrated, and applied to 10 mL of fresh Ni-NTA resin washed with reverse nickel buffer 1 (25 mM Tris pH 7.5, 250 mM NaCl, 20 mM imidazole, and 5 mM BME). The protein was allowed to pass over the nickel resin 3 times, and then washed extensively with reverse nickel buffer 1 until no more protein was washed off the resin as monitored by Bradford Reagent. The protein was then concentrated and added to a Superdex S200 10/300 gel filtration column in a final buffer of 20 mM HEPES pH 7.0, 150 mM NaCl, 5 mM BME. Protein was concentrated to ∼20 mg/mL (∼650 µM) and dialyzed into dialysis buffer 2 (20 mM HEPES pH 7.0, 150 mM NaCl, 3.25 mM DTT) overnight. The next morning 1M CuCl_2_ was added to 650 µM of CCS to a final concentration of 585 µM of CuCl_2_ and mixed until dissolved. The protein loaded with Cu was then flash frozen in liquid nitrogen and stored in −80 °C freezer for future use.

#### Proteins used for surface plasmon resonance

##### T7-HIS-ATOX1

T7-HIS-ATOX1 was expressed in BL21 (DE3) cells in LB with carbenicillin (100 µg/mL, Sigma Aldrich) and when reached the exponential growth, IPTG (Sigma Aldrich) was added. Protease inhibition cocktail (Roche) was added to prevent protein degradation and cells were lysed by sonication. Nucleic acids digestion was performed with universal nuclease (Pierce). The proteins were then purified using an ÄKTA Purifier (GE Healthcare). First, the protein was purified with a HisTrap FF Ni-NTA column (GE Healthcare), using a gradient of imidazole (5 mM – 1 M). Second, further purification was done with a Q Sepharose Fast Flow column (GE Healthcare) and a gradient of NaCl (50 mM – 500 mM). Finally, the last purification step was size exclusion using a HiLoad 16/600 Superdex 75 pg column (GE Healthcare). The protein was concentrated with a 3 kDa Amicon Ultra Centrifugal filter (Sigma-Aldrich).

##### HIS-CCS

HIS-CCS were purchased from Protein Expression Platform at UmeÅ University.

##### HIS-PARVALBUMIN

HIS-PARVALBUMIN purification was previously described(Werner et al., 2018).

##### Proteins used for electrophoretic mobility shift assays (EMSAs) and size exclusion chromatography (SEC)

*8XHIS-CCS* and *6XHIS-MEK1.* 8XHIS-CCS and 6XHIS-MEK1 proteins were expressed in BL21 (DE3) pLysS cells in LB media at 37 °C until reaching an OD600 of 0.6 to 0.8. and then induced with 1 mM IPTG for an additional 4 hours before being harvested. 8XHIS-CCS protein was purified using a HisTrap HP Ni affinity column (Amersham Biosciences) using buffer A (20 mM Tris, pH 8, 300 mM NaCl, and 2 mM DTT) and buffer B (20 mM Tris, pH 8, 300 mM NaCl, 2 mM TCEP, and 1M imidazole). The column was washed with 2% buffer B for 10 bed volume and Ccs eluted with a gradient from 2 to 100% in 80 ml. The 8XHIS-tag was removed from the proteins using TEV protease produced in-house and engineered to contain its own non-cleavable 8XHIS-tag. After digestion overnight at room temperature, the cleaved HIS-tag and TEV protease were removed from the sample by another pass through the nickel column. 6XHIS-MEK1 proteins were purified using buffer A (50 mM HEPES, pH 7.5, 150 mM NaCl, and 1 mM TCEP) and buffer B (50 mM HEPES, pH 7.5, 150 mM NaCl, 1mM TCEP, and 1M imidazole) as described above.

### X-ray Crystallography

Adenylyl-imidodiphosphate (AMP-PNP) and MgCl_2_ were both added to 10 mg/mL MEK1^45-393/Δ264-307/SVQSDI^ to final concentrations of 2 mM. Crystal trays were then set up screening around the crystal condition of 200 mM ammonium fluoride, 18% PEG3350, 4% glycerol, and 10 mM DTT using the hanging drop vapor diffusion method at 4 °C, adapted from a crystal condition used in the literature(Meier et al., 2012). Crystals formed within 2 days and were soaked in mother liquor supplemented with 20% glycerol and either 0 µM or 250 µM of CuCl_2_ and then flash frozen in liquid nitrogen. X-ray diffraction data were collected at a wavelength of 1.37 Å at the Advanced Photon Source (beamline 24-ID-C) for MEK1^45-393/ Δ264-307/SVQSDI^ crystals either unbound or bound with Cu. Both structures were processed using HKL2000 and the anomalous flag was checked for the Cu bound structure.

Both crystal structures were determined by molecular replacement in PHENIX using phaser(Adams et al., 2010; McCoy et al., 2007). The MEK1^45-393/ Δ264-307/SVQSDI^ that was not soaked with CuCl_2_ was determined using PDB 3SLS(Meier et al., 2012) as a search model in phaser and the MEK1^45-393/ Δ264-307/SVQSDI^ soaked with CuCl_2_ was determined by using the Cu-unbound structure as a search model. Molecular replacement search models had waters, metals, and ligands (AMP-PNP or inhibitors) removed from them. For the Cu-unbound structure, the activation segment (residues 210-240) as well as the αC-helix (residues 100-122) were included in the search model but deleted during refinement and rebuilt manually after PHENIX refinement. When identifying the Cu binding site using the anomalous absorption edge of Cu, two additional potential sites were identified near Met187 in chain A and Cys121 in chain B. These signals were much weaker than the main binding site (disappearing at 5.00 and 3.00 sigma, respectively, compared to the ∼13-15 sigma of the main Cu binding site in chains A and B). Due to the ambiguity of these additional potential Cu sites, they were not included in the current model. Model building and refinement were performed using Coot and PHENIX(Afonine et al., 2012; Emsley et al., 2010). For both structures NCS was used, as multiple MEK subunits were present in the asymmetric unit. Anomalous maps were generated by using the “Calculate Maps” tool in PHENIX(Adams et al., 2010). Simulated Annealing omit maps were generated by taking the finished model, deleting the Cu molecule as well as interacting residues (H188, C207, D208,) and running PHENIX refine while checking simulated annealing during the refinement(Afonine et al., 2012). Table 1 statistics were generated using PHENIX validation tools(Chen et al., 2010).

### *In Vitro* Kinase Assay

Activity and activation of MEK1^WT^, MEK1^H188A^, MEK1^H188F^, MEK1^H188M^ MEK1^C207A^, and MEK1^C207S^, by Cu chaperones ATOX1 or CCS were assessed using an ELISA assay adapted from a previously established assay(Binsted and Hasnain, 1996; Grasso et al., 2016). Briefly, GST-ERK2^K54R^ fusion protein was diluted to a final concentration of 5.33 µM in Tris-buffered saline treated with 0.05% Tween-20 (TBST) and added to a glutathione-coated 96 well plate (Pierce, #15240) and incubated at room temperature for 1 hour with shaking. Purified and untagged full-length MEK1^WT^ and mutants were diluted to 400 nM concentrations in kinase buffer 1 (50 mM HEPES pH 7.0 and 50 mM NaCl) and 3 µL of various concentrations of CCS were added to 100 µL of diluted MEK in a 96 well “V” bottom plate (Corning, #2897) to final concentrations ranging from 1.25 µM to 10 µM. The MEK1/ATOX1 or MEK1/CCS mixture was then incubated for 1 hour at room temperature. The glutathione coated plates were then washed extensively with TBST and 50 µL of the MEK1 ATOX or MEK1/CCS mixture was added to the ERK2^K54R^-bound plates along with 50 µL of 200 µM ATP in ATP dilution buffer (50 mM HEPES pH 7.0, 200 mM NaCl, and 20 mM MgCl_2_) bring the final reaction concentrations to 200 nM MEK and 100 µM ATP. The plate was then mixed at room temperature for 5 minutes and then left to incubate in a 37 °C incubator for 30 minutes. The reaction was then washed from the plate and the plate was extensively washed with TBST to quench the reaction. A 1:5000 dilution of rabbit anti-phospho(Thr202/Tyr204)-ERK1/2 (1:1000; Cell Signaling Technology (CST), 9101) in TBST treated with 0.5% BSA was added to the plate and incubated for 1 hour at room temperature with shaking. The plate was then treated to multiple TBST washes and then incubated with a 1:5000 dilution of secondary antibody (goat anti-rabbit IgG (H+L)-HRP (BioRad)) in TBST treated with 0.5% BSA for 1 hour with shaking at room temperature. The plate was again washed with multiple TBST washes and Supersignal ELISA Pico Chemiluminescent Substrate (Pierce, #37069) was added. The plate was read on a Promega GloMAX 96 Microplate Luminometer. Statistical analysis of P-ERK2 luminescence units was analyzed using a one-way ANOVA followed by a Dunnett’s multi-comparisons test or a two-way ANOVA followed by a Tukey’s multi-comparisons test in Prism 7 (GraphPad).

### EPR, XAS, and EXAFS

Samples were dialyzed against 50 mM HEPES containing 150 mM NaCl where necessary to remove BME prior to Cu(II) reconstitution. Cu(II) sulfate was added via syringe pump at a rate of 15 µL per hr during which time a faint precipitate was observed which was more apparent after one mole-equivalent of Cu(II) had been added. In subsequent experiments, Cu(II) addition was terminated at a mole ratio of ∼1.2 Cu/P. The sample was centrifuged to remove any remaining precipitate and passed through two spin desalting columns (ThermoFisher) to remove unbound metal ions. Cu concentrations were measured after dilution on a Perkin-Elmer Optima 2000 ICP-OES instrument. Protein was measured by UV/vis spectrometry using an extinction at 280 nm of 24785 M^−1^cm^−1^.

#### Collection and Analysis of EPR Spectra

Quantitative X-band (9.4 GHz) EPR spectra were recorded on a Bruker Elexsys E500 spectrometer. Temperature control was provided by a continuous nitrogen flow cryostat system, in which the temperature was monitored with Bruker W1100321 thermocouple probe. Frozen Cu(II) reconstituted MEK1 samples at concentrations between 100 – 300 µM in 50 mM HEPES, 150 mM NaCl buffer at pH 7.0 (with no cryoprotectant) were measured in 4 mm quartz tubes at 90 – 110 °K under non-saturating power conditions. Experimental conditions were as follows: microwave frequency 9.678 Ghz; modulation frequency 100 KHz; modulation amplitude 10 G, microwave power 20 mW; field center 3100; field width 1000; time constant 40.96 ms; conversion time 81.92 ms, sweep time 83.89 s.

#### Sample Preparation for XAS

The oxidized protein samples were prepared by concentration to approximately 300 µM. Reduced protein samples were prepared from the Cu(II)-reconstituted samples under anaerobic conditions by addition of a five-fold excess of buffered ascorbate.

#### Collection and Analysis of XAS Data

Samples were measured as aqueous glasses in 20 percent ethylene glycol at 10 K. Cu K-edge (8.9 keV) extended X-ray absorption fine structure (EXAFS) and X-ray absorption near edge structure (XANES) data were collected at the Stanford Synchrotron Radiation Lightsource operating at 3 GeV with currents close to 500 mA maintained by continuous top-off. MEK1^WT^ spectra were measured on beam-line 9-3 using a Si[220] monochromator and a Rh-coated mirror upstream with 12.5 keV energy cutoff in order to reject harmonics. Data were collected in fluorescence mode using a high-count rate Canberra 100-element (BL 9.3) Ge array detector with maximum count rates per array element less than 120 kHz. A nickel oxide filter and Soller slit assembly inserted in front of the detector was used to reduce elastic scattering relative to the Cu Kα fluorescence. Four to six scans of a sample containing only buffer were averaged and subtracted from the averaged data for each protein sample to remove the Ni Kβ fluorescence and produce a flat pre-edge baseline. Data reduction and background subtractions were performed using the program modules of EXAFSPAK (https://www-ssrl.slac.stanford.edu/exafspak.html). Output from each detector channel was inspected for glitches and dropouts before inclusion in the final average. Spectral simulation was carried out using EXCURVE version 9.2 as described previously(Chacón and Blackburn, 2012; Chacón et al., 2014). Simulations of the EXAFS data used a mixed-shell model consisting of imidazole from histidine residues, S from cysteine/methionine coordination, and additional shells (O/N or halide) as necessary. The threshold energy, E0, was chosen at 8985 eV. Refinement of structural parameters included distances (R), coordination numbers (N), and Debye-Waller factors (2σ^2^), and included multiple scattering contributions from outer-shell atoms of imidazole rings.

### BCA Cu(I) affinity

MEK was Cu loaded with CuSO_4_ as described previously with any unbound metal removed with extensive washing with metal-free buffer. ICP-MS was used to determine the concentration of Cu present. Samples for ICP-MS were prepared with 1% HNO_3_. Cu binding experiments were conducted in a nitrogen-purged anaerobic glovebox. Using BCA, the Cu(I) affinity of wild-type MEK1 and MEK1^C207A^ were determined to be 9.24 ± 3.34 ^E-16^ M and E-11 M, respectively. 40 µM of Cu-bound MEK1 or MEK1^C207A^ were incubated overnight at pH 7.4 with 1 mM or 240 µM of BCA and measured spectroscopically using a Cary 300 UV-Vis Spectrophotometer (Agilent). The measured absorbance was used to calculate affinity. All the Cu was removed. The concentration of BCA was lowered to 80 µM which is the lowest usable concentration, and all the Cu again was removed. The Cu(I) affinity for MEK1 was unable to be determined. The lowest affinity range that can be reported using BCA is E-11 M, so the MEK1^C207A^ affinity must be lower than this. All assays were conducted in triplicate, and calculated using the formation constant β_2_=10^17.2^M^−2^ for BCA(Xiao et al., 2011).

### Immunoblot analysis

HEK-HT parental cells or those stably expressing pSMARTvector constructs were treated as indicated and washed with cold PBS and lysed with cold RIPA buffer containing 1X EDTA-free Halt™ protease and phosphatase inhibitor cocktail halt protease and phosphatase inhibitors (Thermo Scientific). Protein concentration was determined by BCA Protein Assay (Pierce) using BSA as a standard. Equal amount of lysates were resolved by SDS-PAGE using standard techniques, and protein was detected with the following primary antibodies: mouse anti-ATP7A (1:1000; 376467, Santa Cruz), mouse anti-β-ACTIN (1:10000; 3700, Cell Signaling), rabbit anti-biotin (1:1000; 5597, Cell Signaling), mouse anti-CCS (1:1000; 55561, Santa Cruz), rabbit anti-phospho(Thr202/Tyr204)-ERK1/2 (1:1000; 9101, Cell Signaling), mouse anti-ERK1/2 (1:1000; 9107, Cell Signaling), rabbit anti-phospho(Ser217/221)-MEK1/2 (1:1000; 9154S, Cell Signaling), mouse anti-MEK1/2 (1:1000; 4694S, Cell Signaling), mouse anti-MEK1 (1:1000; 2352, Cell Signaling), mouse anti-MYC (1:1000; 227S, Cell Signaling), and mouse anti-SOD1 (1:1000; MA1-105, ThermoFisher Scientific) followed by detection with one of the horseradish peroxidase conjugated secondary antibodies: goat anti-rabbit IgG (1:2000; 7074, Cell Signaling) or goat anti-mouse (1:2000; 7076, Cell Signaling) using SignalFire (Cell Signaling) or SignalFire Elite ECL (Cell Signaling) detection reagents. The fold change in the ratio of phosphorylated protein to total protein was measured in Image Studio Lite (LI-COR Biosciences) software by boxing each band using the rectangular selection tool and calculating the signal of the band in pixels. The signal of the phosphorylated protein band in pixels was normalized to the signal of the total protein band in pixels. The average fold change is shown in figures. For BirA proximity-dependent biotin labeling studies, HEK-HT cells stably expressing MYC-BirA, MYC-BirA-ATOX1, or MYC-BirA-CCS were treated with or without biotin for 24 hours as previously described and biotinylated proteins were pulled down with Dynabeads MyOne Streptavidin C1 (Life Technologies, #65002) as previously described.

### Surface plasmon resonance

The interaction between MEK1 and HIS-ATOX1 or HIS-CCS was examined using a Biacore X100 surface plasmon resonance instrument (GE Healthcare). MEK1 was covalently coupled to a CM5 chip (GE Healthcare) using the amine coupling kit according to manufacturer’s instructions (GE healthcare). Briefly, the surface was activated using 0.05 M N-hydroxysuccinimide (NHS) and 0.4 M 1-ethyl-3-(−3-dimethylaminopropyl) carbodiimide hydrochloride (EDC) prior to injection of MEK1 (20 µg/mL in sodium acetate buffer pH 4.5) until 2000 RU (∼2 ng/mm2) was reached and, lastly, 1M ethanolamine was injected for deactivation of the surface. The interaction was measured at 25 °C in HBS-P+ running buffer containing 10 mM HEPES, 150 mM NaCl, and 0.001% P20 detergent at pH 7.4 (GE Healthcare). Increasing concentrations (2x increase for every injection) of HIS-ATOX1 or HIS-CCS were flown over the surface in a single cycle sequence without regeneration steps between injections. At the end of the cycle, regeneration was done by injection of 50 mM NaOH (GE Healthcare). The samples of HIS-ATOX1 and HIS-CCS were prepared with DTT (Sigma Aldrich) with a molar ratio of 5:1 (DTT:protein) and for experiments conducted with Cu (CuCl_2_ from Sigma Aldrich), with a molar ratio of 0.9:1 CuCl_2_:protein. The dissociation constant, K_d_, was determined with the BIAevaluation software using the 1:1 binding model. To confirm that the HIS-tag was not significant for the binding, an equivalent experiment using HIS-PARVALBUMIN was performed.

### Size exclusion chromatography

Size-exclusion chromatography (SEC) of MEK1 and CCS were conducted to determine elution volume using Superdex 75 Increase 10/300 GL (GE Healthcare). Cu(I)-CCS was mixed in equal ratio with MEK1 and the mixture subjected to SEC. The flow rate of each purification was 0.5 ml/min using 50 mM HEPES, pH 7.5, 150 mM NaCl, and 1 mM TCEP buffer. ICP-MS determined the metal content of proteins prior to experimental set-up.

### Electrophoretic mobility shift assays

Metal content of proteins was determined by ICP-MS prior to experimental set-up. Binding assays were performed in triplicate at pH 7.5 in 50 mM HEPES, 150 mM NaCl, and 1 mM TCEP. Assays were conducted using apo-CCS and Cu(I)-CCS with CCS in excess and MEK1 constant. Experimental concentrations were chosen based on the affinity determined previously by SPR at 8 µM. Protein reactions were allowed to incubate 20 minutes at room temperature prior to native gel analysis. 10% native gels were loaded with 10ul of each reaction with 2ul of 50% glycerol added. Gels were run at 150V for 60 minutes and then stained with Coomassie blue for 20 minutes. Gels were de-stained overnight with agitation and visualized using the ChemiDoc MP system (BioRad).

## Supplemental Information

**Figure S1.**
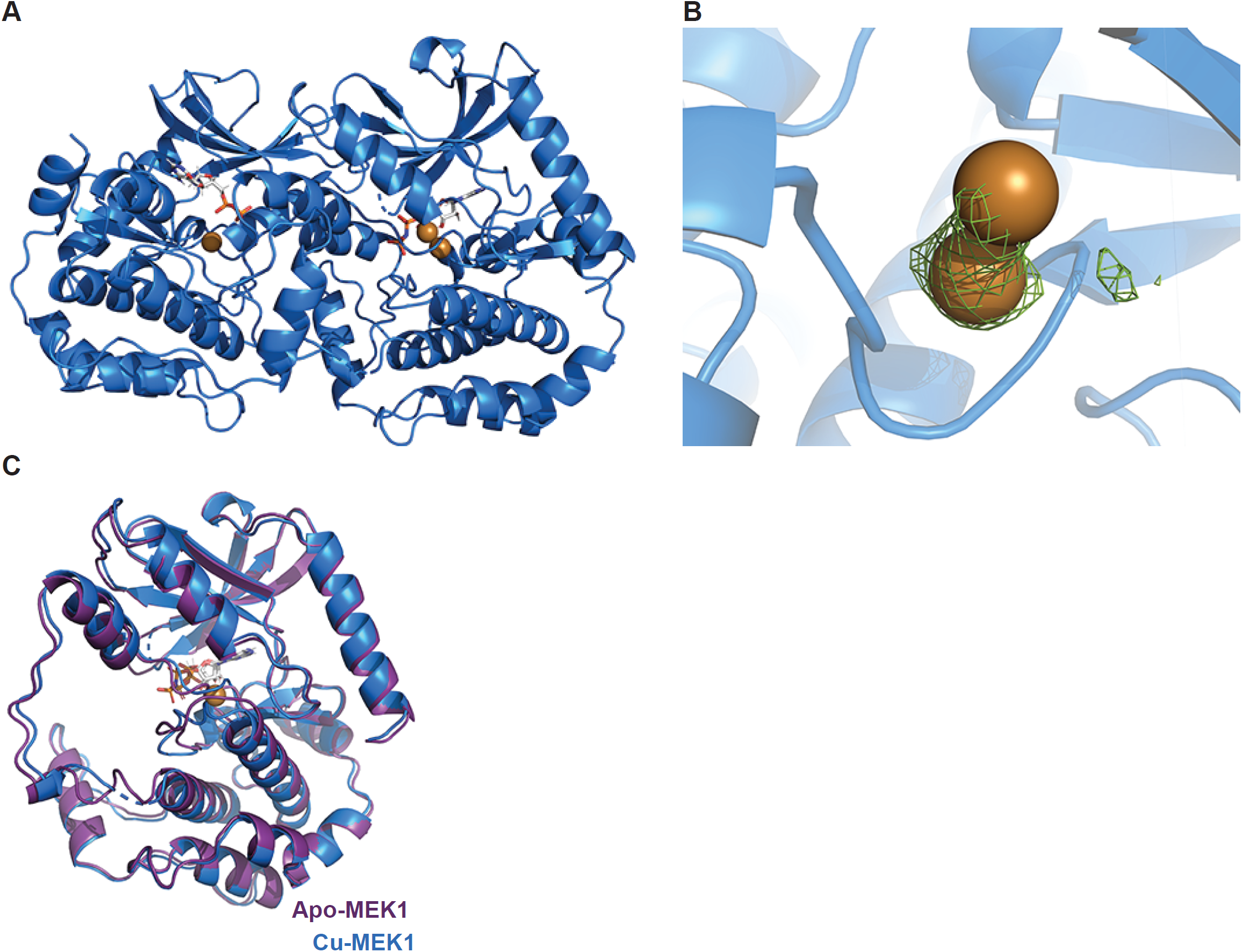
Alternate conformation of Cu atom in MEK1 X-ray crystal structure. (A) Crystallographic dimer of MEK1^45-393/SVQSDI^ (marine blue) bound with Cu ions (brown) and AMP-PNP (grey). Chain A demonstrates 2 conformations of Cu built in to the model. (B) Close-up of Cu (brown) in both conformations and its anomalous map (lime green, contoured at 3.0 sigma) bound to MEK1 (marine blue). The major conformation, which makes contacts with sidechains of H188 and C207, and the main chain of D208, has 80% occupancy while the second Cu atom has 20% occupancy. (C) Alignment of crystal structures of apo MEK1 MEK1^45-393/SVGSDI^ (purple) bound with AMP-PNP (grey) and MEK1^45-393/SVGSDI^ (marine blue) bound with Cu ions (brown) and AMP-PNP (grey).

**Figure S2.**
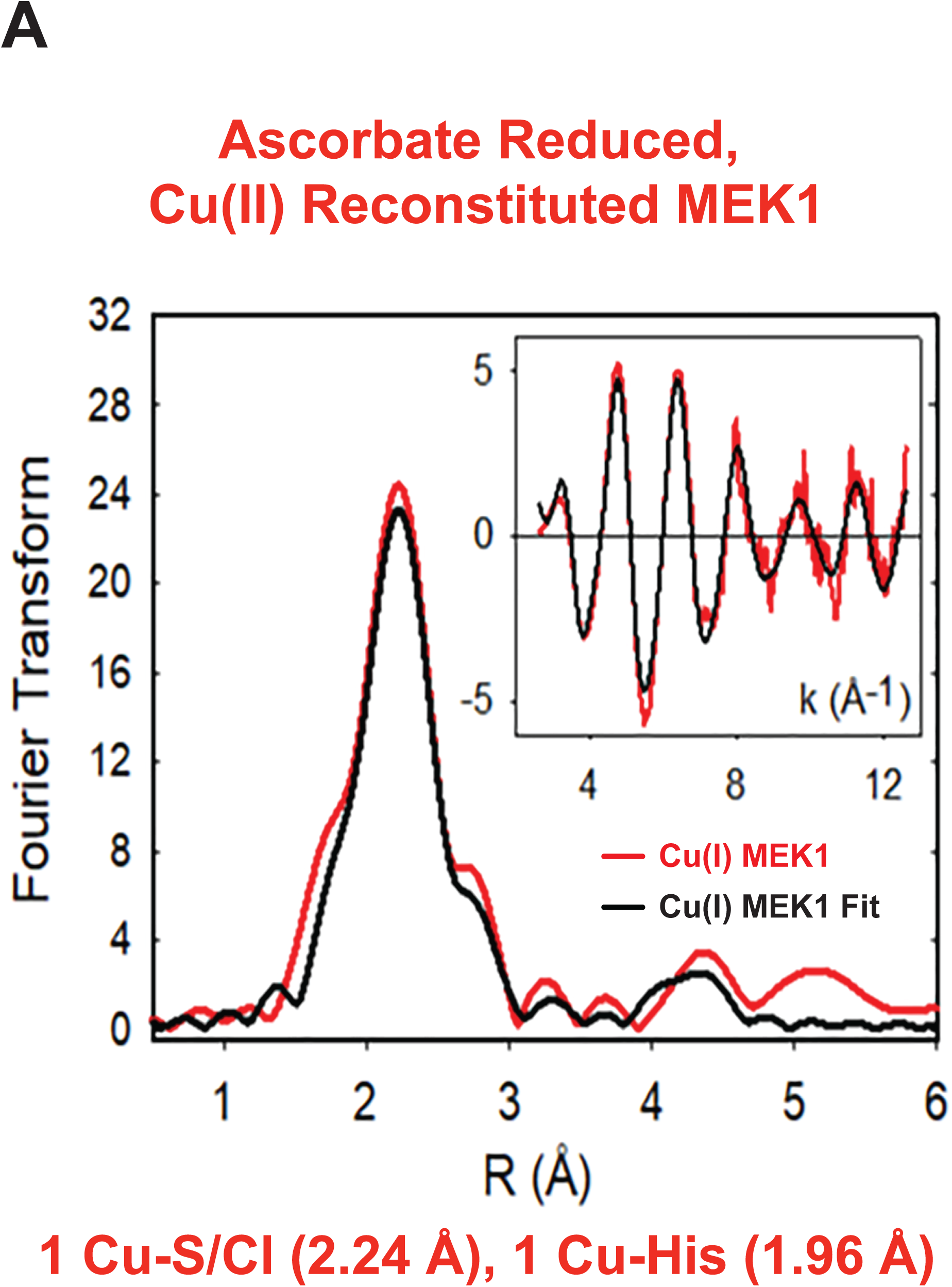
MEK1 binds Cu(I) via histidine and cysteine ligands. (A) Fourier transform and EXAFS (inset) of experimental ascorbate reduced, Cu(II) reconstituted MEK1 in NaBr (red line) and its fit (black line). n=1 technical replicate.

**Figure S3.**
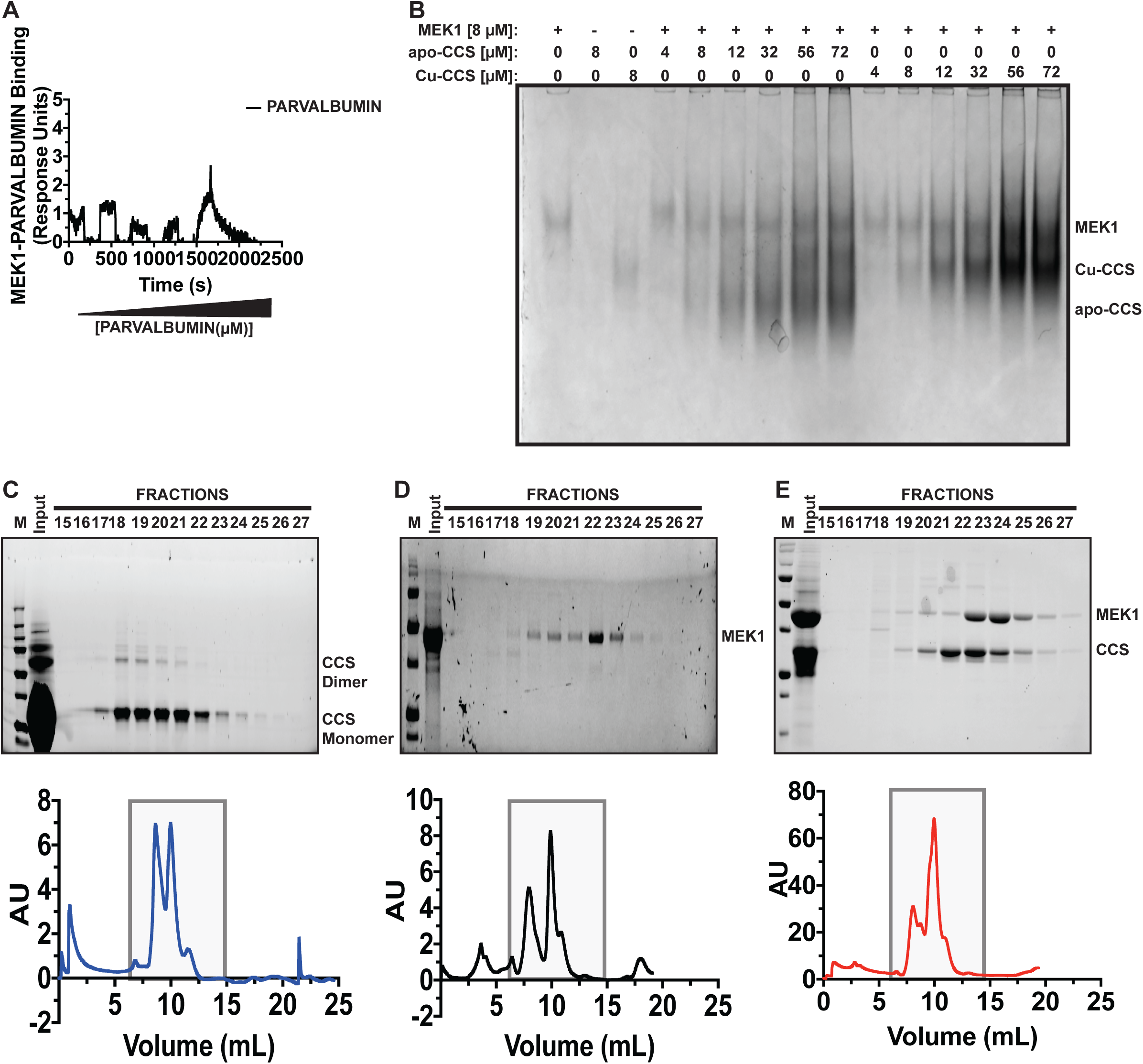
MEK1-CCS fail to form a constitutive complex. (A) Single-cycle SPR experiments of HIS-tagged PARVALBUMIN to MEK1 in which increasing concentrations of PARAVALBUMIN (2.5-40µM, black line) was injected onto immobilized MEK1. n=3 independent experiments. (B) Electrophoretic mobility shift assays (EMSAs) of increasing concentrations of apo-CCS or Cu reconstituted CCS (Cu-CCS) with increasing concentrations of MEK1. (C-E) Size exclusion chromatography runs of (C), Cu-CCS (blue line), (D), MEK1 (black line), or (E), Cu-CCS and MEK1 (red line) and protein detection in indicated fractions detected by Coomassie Brilliant Blue.

